# The oxygen tolerant reductive glycine pathway in eukaryotes – a native methanol, formate and CO_2_ assimilation pathway in the yeast *Komagataella phaffii*

**DOI:** 10.1101/2022.09.01.506198

**Authors:** Bernd M. Mitic, Christina Troyer, Stephan Hann, Diethard Mattanovich

## Abstract

The current climate change is mainly driven by excessive anthropogenic CO_2_ emissions. As industrial bioprocesses depend mostly on food competing organic feedstocks or fossil raw materials, we regard CO_2_ co-assimilation or the use of CO_2_-derived methanol or formate as carbon source as pathbreaking contribution to the solution of this global problem. The number of industrially relevant microorganisms that can use these two carbon sources is limited, and even less can concurrently co-assimilate CO_2_. Hence, we searched for alternative native methanol and native formate assimilation pathways which co-assimilate CO_2_ in the industrially relevant methylotrophic yeast *Komagataella phaffii* (*Pichia pastoris*). Using ^13^C-tracer-based metabolomics techniques and metabolic engineering approaches we discovered and confirmed a natively active pathway that can perform all three assimilations: the oxygen tolerant reductive glycine pathway. This finding paves the way towards metabolic engineering of formate and CO_2_ utilisation for the production of proteins, biomass or chemicals in yeast.

## Introduction

The combustion of fossil fuels and the associated increase of atmospheric CO_2_ concentration is the primary reason of anthropogenic climate change^1^. Regarding this global problem, CO_2_-fixation is of utmost importance. The use of CO_2_ derived renewable and green feedstocks as carbon sources for bioprocesses has increased in importance over the last years not only due to climate change concerns, but also due to the problems emerging from food competing bio-industrial feedstock, e.g. glucose. Green, non food competing carbon sources such as methanol (MeOH) and formate (FA) can be produced electrochemically from CO_2_^2,3^. Still, the number of organisms which can produce biomass, proteins or chemicals on these green carbon sources is limited^4^.

*Komagataella phaffii* (also known as *Pichia pastoris*) is a methylotrophic yeast^5,6^, which is industrially used for heterologous protein production of enzymes and biopharmaceuticals and is well-known for its methanol inducible alcohol oxidase promotor system^7–10^. Synthetic biology tools, including CRISPR/Cas9, make metabolic engineering easily applicable to this methylotrophic host^11–14^. This was recently demonstrated by changing the heterotrophic organism in an auxotroph which can grow on CO_2_^15^. The main methanol assimilation pathway, the xylulose 5-phosphate pathway (XuMP), is well understood and has been investigated in detail^16^. Hitherto unknown metabolic routes were recently found in *K. phaffii*. These routes are active even though their flux is too low to support growth^17^. These findings motivated us to seek for other hidden pathways, focusing on methanol, formate and CO_2_ assimilation routes which are natively active. In nature several methanol fixation pathways are present and active. They are differentiated into formaldehyde- and formate-fixing pathways (see Fig. 1). Like the XuMP the ribulose 5-phosphate pathway (RuMP) of *Bacillus methanolicus* is a formaldehyde-fixing pathway (Fig. 1, red & orange). It uses the pentose phosphate pathway (PPP) to recycle a sugar-phosphate and produce glyceraldehyde phosphate (GAP), which is used for biomass formation^18,19^. The enzymes 3-hexulose 6-phosphate synthase (Hps) and phosphohexose isomerase (Phi) are of central importance in the RuMP as shown in engineered *Escherichia coli* for formaldehyde consumption and growth^20^ and for methanol consumption^21^. After adaptive laboratory evolution also *Saccharomyces cerevisae* can grow on methanol through a formaldehyde fixing pathway which leads through the PPP^22^. In the serine cycle pathway of *Methylorubrum extorquens* (formerly *Methylobacterium extorquens*^23^) methanol is dissimilated to formaldehyde and further to formate in a formate-fixing pathway, before entering the enzymatic tetrahydrofolate (THF) pathway, leading to methylene-THF^24,25^. Besides this formate-fixing pathway it may, to a minor extent, also be formaldehyde-fixing due to a spontaneous *in vivo* condensation reaction of THF with formaldehyde forming methylene-THF^24,26^(see Fig. 1, cyan). In the serine cycle itself, the carbon originating from methanol or formate is condensed with a second carbon from CO_2_ ending up in acetyl-CoA, which is used for all biomass formation ^24,27^ (see Fig. 1, blue reactions in the cycle). Therefore, the serine cycle is a CO_2_-fixing pathway. An engineered version of the serine cycle via pyruvate following a different CO_2_-fixation reaction was demonstrated in *E. coli*^28^ (see Fig. 1, straight blue reactions).

**Figure 1:**
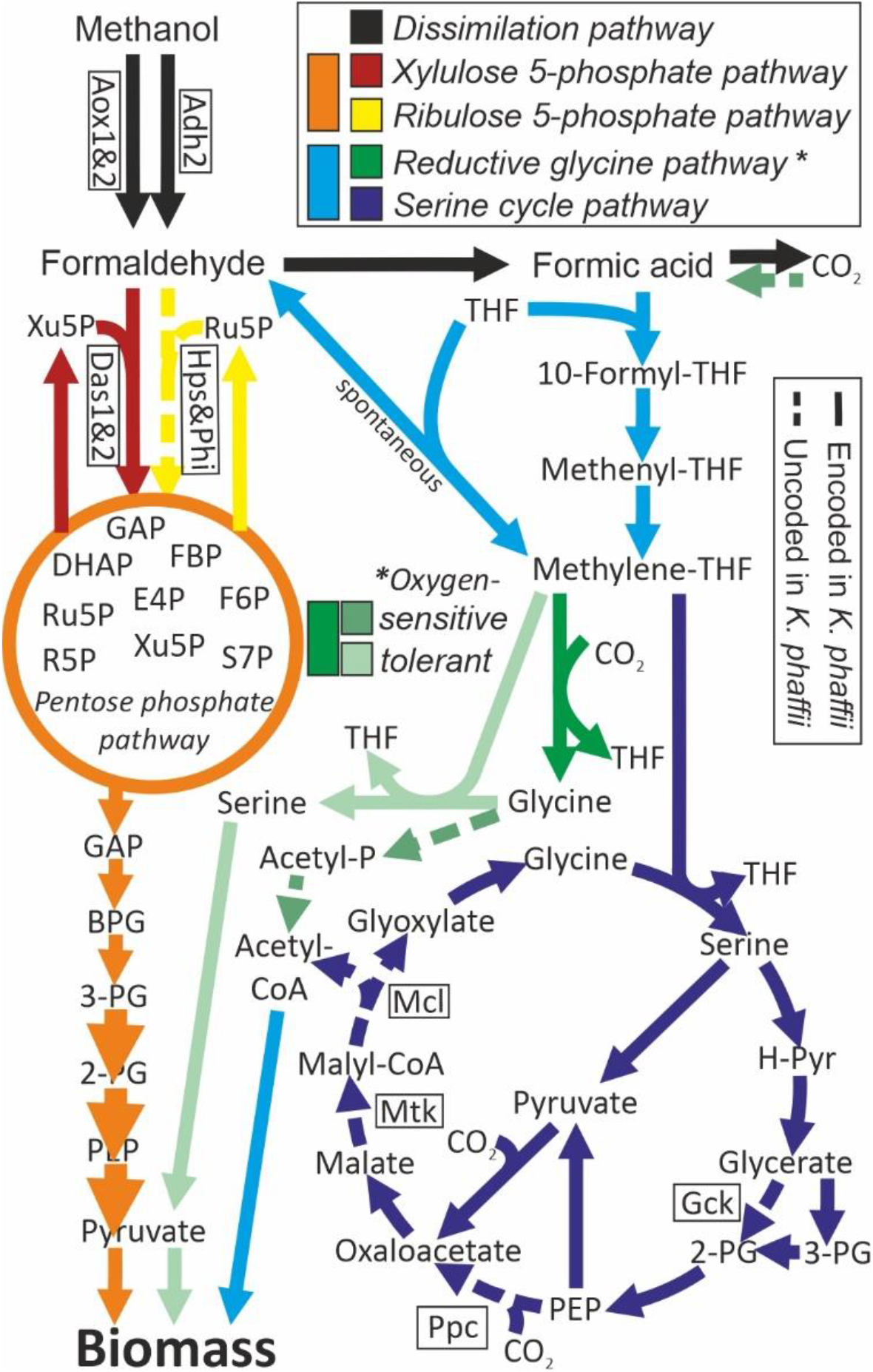
Scheme describing methanol and formate assimilation pathways. The xylulose 5-phosphate pathway, the native and main methanol assimilation pathway in *K. phaffii*, and the ribulose 5-phosphate pathway (*B. methanolicus*) are methanol assimilation pathways and fix formaldehyde to produce glyceraldehyde phosphate through pentose phosphate pathway reactions, serving as metabolite for all biomass production. The reductive glycine pathway and the serine cycle are methanol and formate assimilation pathways, as methanol gets dissimilated and formate is fixed in the tetrahydrofolate cycle and both co-assimilate CO_2_. The O_2_-tolerant reductive glycine pathway leads to pyruvate, while the O_2_-sensitive reductive glycine pathway (*D. desulfuricans*) and the serine cycle (*M. extroquens*) end up in acetyl-CoA as metabolite for biomass production. Reactions shown focus on carbon reactions only, for other involved compounds, enzyme and gene names see Supplementary information Fig. S1&2. Dashed reactions are not encoded in *K. phaffii*. Abbreviations: THF – tetrahydrofolate, R5P – ribose 5-phosphate, S7P – sedoheptulose 7-phosphate, E4P – erythrose 4-phosphate, Xu5P – xylulose 5-phosphate, Ru5P – ribulose 5-phosphate, FBP – fructose bis-phosphate, DHAP – dihydroxyacetone phosphate, GAP – glyceraldehyde phosphate, BPG – bis-phosphoglycerate, 2-& 3-PG – 2-& 3-phosphoglycerate, PEP – phosphoenolpyruvate, H-Pyr – hydroxyl-pyruvate, Acetyl-P – acetyl-phosphate, Adh – alcohol dehydrogenase, Aox – alcohol oxidase, Das – dihydroxyacetone synthase, Hps – 3-hexulose-6-phosphate synthase, Phi – phosphohexose isomerase, Gck – glycerate 2-kinase, Pdc – phosphoenolpyruvate carboxylase, Mtk – malate-CoA ligase, Mcl – malyl-CoA lyase

The oxygen-sensitive reductive glycine pathway is the native carbon and CO_2_-fixation pathway of the anaerobic bacterium *Desulfovibrio desulfuricans*^29^. CO_2_ is reduced to formate by an oxygen-sensitive enzyme leading to methylene-THF as in the serine cycle. In the reaction defining the general reductive glycine pathway, a second CO_2_ molecule is fixed and enters the *de novo* synthesized amino acid glycine via the glycine cleavage system. Glycine is metabolized to acetyl-phosphate by another oxygen-sensitive reaction before ending up in acetyl-CoA, which can either be directly used for biomass production or partially fixes a third carbon dioxide molecule forming pyruvate^29^. The reductive glycine pathway from formate to glycine was overexpressed in a glycine auxotrophic *S. cerevisiae* strain and led to growth on glucose, formate and CO_2_, which proved the reversibility of the glycine cleavage system in yeasts^30^. In an alternative route, natively only known on a genomic level, glycine reacts with a second molecule of methylene-THF to serine, which is then deaminated to pyruvate, which serves as a precursor for all biomass formation. This so-called oxygen tolerant reductive glycine pathway is known from metagenomic analysis as pure CO_2_-fixation route^31^ and was also bioinformatically designed and integrated as a synthetic pathway in *E. coli* where it supported growth on formate and methanol in combination with CO_2_ ^32–34^ or on formate and glycine^35^. Still, a proof of native metabolic activity of the oxygen tolerant reductive glycine pathway could not be found in nature for any organism.

Our first objective was to search for alternative methanol assimilation pathways in *K. phaffii*. For this purpose, we used a XuMP knockout strain (*das1 Δdas2Δ*) of *K. phaffii* to trace potential alternative methanol incorporation using ^13^C-methanol^17^. To search for potential formate assimilation routes ^13^C-formate labelling was applied to the wildtype. To trace CO_2_-assimilation we used our established reverse labelling approach^15^. The labelling experiments were evaluated using tailored gas chromatography high resolution mass spectrometry (GC-HRMS) metabolomics methods, in more detail GC-chemical ionisation (CI)-time of flight (TOF) MS^36^ and GC – electron ionisation (EI) TOFMS exploiting the potential of different derivatization methods. We support our findings with specific knockouts and targeted overexpression of metabolic routes to gain insights into natively active metabolic pathways in the eukaryote *K. phaffii*.

Such natively active pathways for methanol and formate assimilation and concurrent CO_2_ assimilation are of importance as they can serve as a platform for sustainable bioprocesses of this industrially relevant yeast in the future.

## Results

### ^13^C-methanol labelling indicates the presence of the reductive glycine pathway *in K. phaffii*

When cultivating the XuMP knockout (DasKO) strain on ^13^C-methanol, the presence of an alternative methanol assimilation pathway became apparent by the increase of the ^13^C content of various metabolites over time (Fig. 2a). Pathway routes were assessed by tracing the relative ^13^C abundance in metabolites as shown in Fig. 2a-c. In these figures the number x of ^13^C-atoms in a metabolite is denoted by ‘‘M+x’
s’, thus specifying the respective isotopologue. An increase of ^13^C-content is always associated with a decrease of the M+0 (^12^C only) isotopologue fraction. This M+0 isotopologue fraction is also indicated as numeric value in the corresponding bar for all forward labelling approaches. Generally, an upstream metabolite of any active pathway contains a higher fraction of ^13^C than the corresponding downstream metabolites. Compounds showing the fastest and highest incorporation of the ^13^C label are indicative for the beginning of an active native pathway. Of all measured metabolites, methionine and serine showed both a strong decrease of the unlabelled fraction M+0 of 6-8% already after 2 hours of labelling and the highest label incorporation at all three sampling points. These data clearly show that methanol is - alternatively to the XuMP pathway - assimilated mainly through the tetrahydrofolate pathway, as this pathway directly produces methionine and serine. Serine showed an increase of the isotopologue M+2 after 24 h, which indicates that a second labelled carbon atom, originating from methylene-THF, is incorporated via glycine. Glycine increased in ^13^C content after 24 h of labelling. The mass spectrometric fragment of serine and aspartate, which is specific for the amino acid backbone (BB) as it contains the C1 and C2 carbons of the amino acids only, showed a very similar labelling pattern as glycine. This data further supports the involvement of glycine and the enzyme serine hydroxymethyltransferase (Shm) in the alternative methanol pathway. After 24 hours an increase of the ^13^C content of aspartate and malate was observed as well. Malate and aspartate are direct downstream metabolites of oxaloacetate, which is produced by carboxylation of pyruvate. The appearance of ^13^C in malate and aspartate is therefore the first evidence that serine is deaminated to pyruvate, serving as the labelled precursor. The labelling pattern of oxaloacetate was inaccessible as its concentration in the extracts was below the method’s limit of detection. Pyruvate was found to be a degradation product of oxaloacetate and phosphoenolpyruvate (PEP), metabolites of two different pathways. Hence this metabolite could not be evaluated. Generally, such metabolic interconversion reactions (non-enzymatic *in vitro* reactions) can falsify results and consequently pathway interpretation. Therefore, we tested critical analytes of the hypothesized pathways in this regard (detailed information in Supplementary information Chapter 1.3.1). If formaldehyde fixing pathways leading through the pentose phosphate pathway (PPP) and glycolysis were active, we would expect a different pattern: the metabolites of these pathways, such as sedoheptulose 7-phosphate (S7P), ribose 5-phosphate (R5P), 2&3-PG, would show an earlier and higher degree of ^13^C labelling than methionine, serine and glycine, as well as the downstream metabolites aspartate, malate and fumarate. This pattern can indeed be seen in the control experiment, a methanol labelling experiment with a non-growing Mut-strain, which still harbours the native xylulose 5-phosphate pathway^17^ (Fig. 2 b). The low degree of ^13^C-labels observed in glycolysis and pentose phosphate pathway metabolites in the DasKO strain can be explained by incorporation of ^13^C from oxaloacetate via gluconeogenesis. The native serine cycle route of *M. extorquens* through glycerate, 2-PG and 3-PG to oxaloacetate (see Fig. 1) is obviously not active in *K. phaffii* as malate and aspartate show higher labelling values than 2-PG, 3-PG and PEP. Still, as reflected by the labelling data, the reactions of the synthetic serine cycle via pyruvate to malate were active. If the reactions resulting in acetyl-CoA were active, we would expect a higher ^13^C content in the tricarboxylic acid (TCA) cycle metabolites isocitrate (I-Cit) and α-ketoglutarate (AKG), which are the first measurable metabolites after the acetyl-CoA fixation point of the TCA cycle. Furthermore, ^13^C labelling degrees decreasing from malate and fumarate to AKG and I-Cit indicate a reversely active TCA cycle.

**Figure 2:**
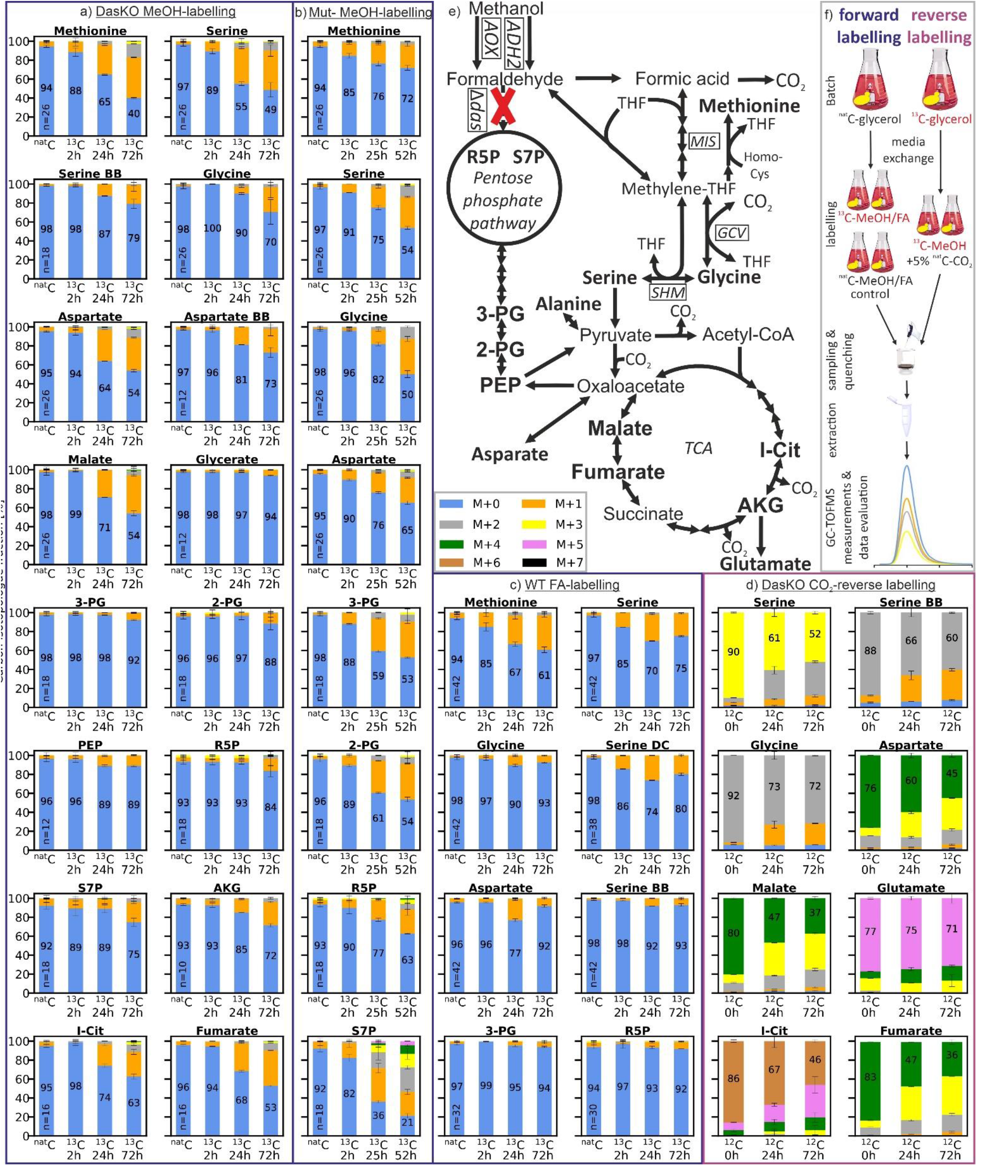
Carbon isotopologue distribution analysis via GC-CI/EI-TOFMS. a-e) *K. phaffii* strains (see Table 1) labelled with different carbon sources (n=2 biological replicates for labelled strains, number of replicates (n) of the ^nat^C controls is indicated in the bar). “BB” in the metabolite name stands for the amino acid backbone, i.e. C1 and C2 only, “DC” stands for the decarboxylated amino acid, i.e. all carbon atoms except C1 (see Supplementary Table S6). The number x in ‘‘M+x’
s’ indicates the number of ^13^C-carbon atoms, thus specifying the respective isotopologue. Pathway routes were assessed by tracing the relative ^13^C abundance in metabolites as shown in a-c for forward labelling experiments. Generally, an upstream metabolite of any active pathway has to contain more ^13^C than the corresponding downstream metabolite. An increase of ^13^C-content results in a decrease of the isotopologue M+0, containing ^12^C only. For all forward labelling approaches, the isotopologue fraction of M+0 is also indicated as number in its corresponding bar, see a-c). For reverse labelling (2d), ^12^C incorporation is traced, therefore any decrease in abundance of isotopologues containing ^13^C indicates CO_2_ incorporation. a) *das1Δdas2Δ* strain labelled with ^13^C-methanol for 72 h; b) *aox1Δaox2Δ* strain labelled with ^13^C-methanol for 52 h; c) wildtype strain labelled with ^13^C-sodium formate for 72 h; d) ^13^C-labeled *das1Δdas2Δ* strain reverse labelled with ^nat^C-CO_2_ (tracing of ^12^C); e) reductive glycine pathway and native xylulose 5-phosphate pathway illustrated down to the TCA cycle, all measured metabolites are highlighted; f) illustration of labelling and reverse labelling workflow. Labelling data of more metabolites see Supplementary Figure S7 a-f; Abbreviations: MeOH – methanol, FA – formic acid, THF – tetrahydrofolate, TCA – tricarboxylic acid cycle, R5P – ribose 5-phosphate, S7P – sedoheptulose 7-phosphate, 2-PG – 2-phosphoglycerate, 3-PG – 3-phosphoglycerate, PEP – phosphoenolpyruvate, AKG – α-ketoglutarate, I-Cit – isocitrate

When performing the methanol labelling while supplying additionally ^nat^C glycine in the medium, the same labelling pattern was observed, however with reduced labelling intensity, which results from the reaction of glycine to methylene-THF leading to a mixed feed of methanol and glycine. This additionally proves the general activity of the glycine cleavage system. (see Supplementary information Fig. S7 e; labelling results for more metabolites than shown in Fig. 2 a&b can be found in Supplementary information Fig. S7 a&b.) Briefly, it can be concluded from the ^13^C methanol labelling data obtained from the DasKO strain that the oxygen tolerant reductive glycine pathway from methanol to oxaloacetate is the main active alternative methanol assimilation pathway in *K. phaffii*.

**Table 1:** Names and genotypes of strains used in this study

| Strain name | Genotype | Source |
| --- | --- | --- |
| **WT** | CBS7435 |  |
| **DasKO** | CBS7435 *das1Δdas2Δ* | This study |
| **GcvKO** | CBS7435 *das1Δdas2Δ gcv1Δgcv2Δ* | This study |
| **ShmKO** | CBS7435 *das1Δdas2Δ shm1Δshm2Δ* | This study |
| **MisKO** | CBS7435 *das1Δdas2Δ mis1-1Δmis1-2&3Δ::loxP-kanMX-loxP* | This study |
| **GcvOE** | CBS7435 *das1Δdas2Δ P<sub>FDH1</sub>GCV1 P<sub>DAS2</sub>GCV2 P<sub>AOX1</sub>LPD1 P<sub>DAS1</sub>GCV3* | This study |
| **OtRedGlyOE** | CBS7435 *das1Δdas2Δ P<sub>FDH1</sub>GCV1 P<sub>DAS2</sub>GCV2 P<sub>AOX1</sub>LPD1 P<sub>DAS1</sub>GCV3*<br>*P<sub>DAS1</sub>MIS1-1 P<sub>AOX1</sub>SHM1 P<sub>DAS2</sub>CHA1 P<sub>DAS1</sub>ADE3 P<sub>AOX1</sub>SHM2* | This study |
| **Mut-** | CBS2612 *aox1Δaox2Δ* | Zavec et al.<br>2021<sup>17</sup> |
| **Das&Aox1KO** | CBS7435 *das1Δdas2Δ aox1Δ* | Gassler et al.<br>2020<sup>15</sup> |

### ^13^C-formate labelling demonstrates the oxygen tolerant reductive glycine pathway as native formate assimilation route in *K. phaffii*

Formate (FA) is an intermediate metabolite of methanol dissimilation. If the oxygen tolerant reductive glycine pathway is the alternative methanol assimilation pathway in *K. phaffii*, also formate can be used as a carbon source for this organism via this pathway. Hence, a ^13^C-formate tracer experiment was designed and performed for further validating the activity of this pathway. The result should assess both, that formate is assimilated, and formaldehyde is generated directly from formate or via the spontaneous degradation of methylene-THF to THF and formaldehyde. As depicted in Fig. 2 c, the formate labelling data of the wildtype strain showed a comparable labelling pattern as the methanol labelling data (Fig. 2 a) for the sample points 2 and 24 hours. Methionine and serine are labelled earliest and most substantially. Glycine, aspartate and the serine backbone, which reflects the carbon atoms in serine stemming from glycine, were significantly labelled at the second sample point (24h) and labelled to a higher degree than the metabolites of glycolysis (3-PG) and the pentose phosphate pathway (R5P). At 72 hours the labelling degree was decreased for all analysed metabolites except methionine, going along with reduced consumption of formate after 24 hours of cultivation as shown by analysis of the culture medium (see Supplementary information Fig. S8). Formate dissimilation provides only half the energy than methanol (one NADH, compared to two from methanol), obviously leading to severe starvation after 24h of cultivation. Starvation induced autophagy is the likely source for a pool of unlabelled metabolites at the late stage of this experiment leading to lower labelling degrees at 72 h in comparison to 24 hours. Formate-labelling of the DasKO strain resulted in similar patterns for all time points and metabolites compared to the wild type strain (see Supplementary information Fig. S7 f). Hence *K. phaffii* can natively assimilate formate through the oxygen tolerant reductive glycine pathway. Any reverse reaction to formaldehyde followed by fixation via the xylulose 5-phosphate pathway was not observed as the wildtype and the DasKO showed similar labelling patterns. Furthermore, the data indicate that ^13^C labelling in the pentose phosphate pathway and in glycolysis, in all experiments, derive from gluconeogenesis starting from oxaloacetate, as postulated in the previous section. (Labelling data of more metabolites: Supplementary information Fig. S7 c)

### CO_2_ reverse labelling proves that methanol and CO_2_ are co-assimilated to pyruvate and further to oxaloacetate

To trace CO_2_ co-assimilation via the reductive glycine pathway, more specifically via the glycine cleavage system, we performed a CO_2_ reverse labelling experiment. Avoiding the usage of ^13^C-CO_2_, we fully labelled the DasKO strain with ^13^C-glycerol followed by reverse labelling with ^nat^C-CO_2_ under ^13^C-methanol addition (see Fig. 2 f). For reverse labelling, ^12^C incorporation is traced, therefore any decrease in abundance of isotopologues containing ^13^C indicates CO_2_ incorporation. The experiment clearly revealed that native CO_2_-fixation pathways are active, because the ^13^C content of the metabolites decreased over cultivation time, i.e. the isotopologue distribution pattern was shifted towards isotopologues with a lower number of ^13^C atoms (Fig. 2 d). Serine and glycine were also reverse labelled, which is further evidence of the activity of the anabolically acting glycine cleavage system leading to CO_2_-fixation. A second carboxylation reaction in this route from pyruvate to oxaloacetate is verified by the fact that aspartate and malate showed most intense reverse labelling. The increase of the isotopologues M+3 and M+2 over time displays the double CO_2_-fixation of this oxygen tolerant reductive glycine pathway to oxaloacetate, both via carboxylation by the glycine cleavage system and via carboxylation by pyruvate carboxylase. If a third native CO_2_-fixation route, the reverse reaction of the TCA cycle, was active, a more substantial increase of ^12^C in isocitrate than in malate and fumarate was expected. As this was not the case, activity of the complete reverse TCA cycle could not be verified. (Labelling data of more metabolites: Supplementary information Fig. S7 d

### The oxygen tolerant reductive glycine pathway is the only alternative methanol, formate and CO_2_ co-fixing pathway encoded in the genome of *K. phaffii*

The xylulose 5-phosphate pathway (XuMP) is the most active methanol assimilation pathway in *K. phaffii* and has been studied in detail^16^. In order to identify alternative methanol assimilation pathways *in silico*, we considered the native formaldehyde fixing dihydroxyacetone synthase (*DAS1, DAS2*) as deleted and compared natively encoded enzymes and resulting pathways to methanol assimilation pathways of other methylotrophic organisms. For the ribulose 5-phosphate pathway (RuMP), only 3-hexulose-6-phosphate synthase (Hps) and phosphohexose isomerase (Phi) are missing in *K. phaffii* (Fig. 1, yellow dotted reaction). Still, we have to consider native but unknown formaldehyde-condensing aldolase activities in *K. phaffii*^37^, such as in the evolved *S. cerevisae* strain mentioned in the introduction^22^. In *K. phaffii*, all formate assimilation pathways are also methanol assimilation pathways, as formate is produced in the dissimilation pathway of methanol (Fig. 1, black reactions). For the native serine cycle, a formate assimilation pathway, four enzymes are not encoded in *K. phaffii* (Fig. 1, blue dotted reactions). Considering the shortcut via pyruvate, only malate-CoA ligase (Mtk) and malyl-CoA lyase (Mcl) are not encoded for acetyl-CoA synthesis. All other necessary enzymes from methanol or formate to malate are encoded (Supplementary information Fig. S2) and active as demonstrated in the labelling experiments.

To find potential additional CO_2_ assimilation pathways *in silico*, we searched for CO_2_ fixing enzymes and pathways inspired by organisms which can grow on CO_2_ as sole carbon source. For the oxygen-sensitive reductive glycine pathway, as found in *D. desulfuricans*, the oxygen-sensitive glycine reductase complex to acetyl-phosphate and the phosphate acetyltransferase or the acetate kinase to acetyl-CoA are not encoded in the genome of *K. phaffii* (see Fig. 1, light green doted reactions). Other native or synthetic methanol or formate assimilation pathways are not probable either due to the need of anaerobic conditions, e.g. the reductive acetyl-CoA pathway, or the lack of many genes, which is the case for the homoserine cycle^38^, the serine-threonine cycle and other synthetic routes proposed by Bar-Even in 2016^39^. The oxygen tolerant reductive glycine pathway is the only pathway of which all enzymes were found to be encoded (see Fig. 1, green reactions to pyruvate; details in Supplementary information Fig. S1). This bioinformatic analysis confirm the results of the labelling experiment and strengthens the hypothesis that the oxygen tolerant reductive glycine pathway is a native, metabolically active methanol, formate and CO_2_ assimilation pathway in *K. phaffii*.

During genome mining we found the cytosolic folate pathway gene MIS (encoding C1 tetrahydrofolate synthase) to be split into 2 genes, leading to a higher expression of the formate-tetrahydrofolate ligase subunit as compared to the methenyltetrahydrofolate cyclohydrolase and dehydrogenase subunits. This split might lead to a higher flux towards 10-formyl-THF formation on methanol, which is for instance needed for purine *de-novo* synthesis (see Supplementary information Chapter 1.1.1).

### The native reductive glycine pathway is too limited to allow growth on methanol or formate combined with CO_2_

Growth experiments were performed to further characterize the strains used in the labelling experiments, to assess native flux limitations of the alternative methanol assimilation pathway in order to support growth, and to investigate the activities of specific parts of the active oxygen tolerant reductive glycine pathway (results in Fig. 3).

**Figure 3:**
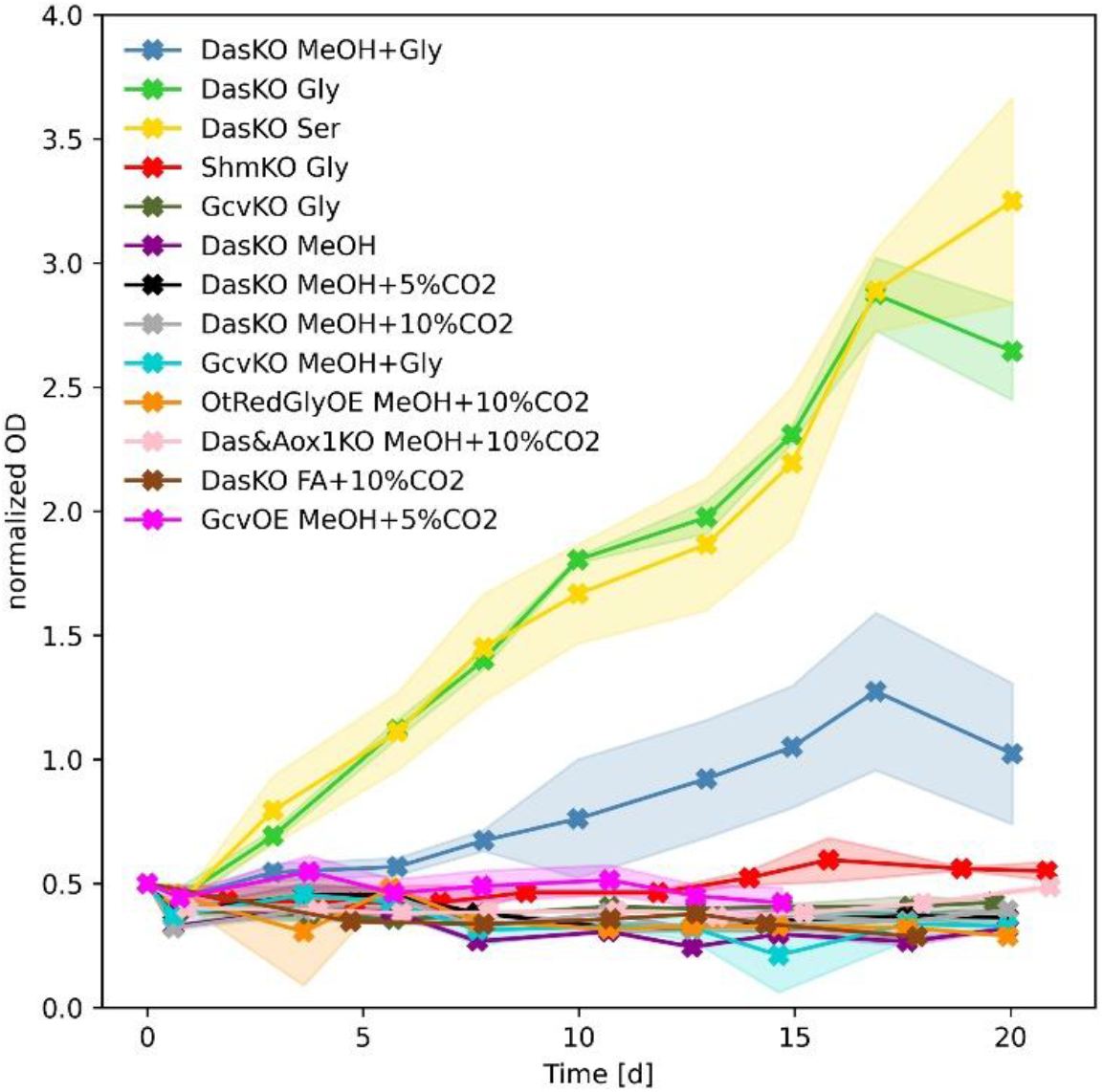
Results of growth analysis. Cultures were started at OD_600_around 0.5, and all data normalized to a starting value of 0.5 for better comparability. All tested strains have at least *DAS1&2* knockouts, some strains have further knockouts or overexpressions as indicated (genotypes are listed in Table 1). Data are average values from biological duplicates with standard deviation indicated in shades. Long-term cultivations (up to 46 days), repeatability studies and further cultivations of overexpression strains are shown in Supplementary information Figure S6.

The xylulose 5-phosphate pathway knockout strain (DasKO) can neither grow on methanol (8 g L^-1^) alone, nor on methanol (8 g L^-1^) or formate (30 mmol L^-1^) at elevated CO_2_ levels of 5% or 10%. Still, the cells consume these carbon sources and formate is even secreted when methanol is dissimilated (see Supplementary information Fig. S8). As formate is produced intracellularly via the methanol dissimilation pathway, all other conditions were tested just on methanol, not on formate. Growth of the DasKO strain could only be observed on methanol when combined with glycine (20 mmol L^-1^) or on glycine or serine only, indicating that the oxygen tolerant reductive glycine pathway starting from glycine ending up in biomass is natively active enough to support growth. The involvement of the glycine cleavage system (Gcv) as a prerequisite for the growth on glycine was affirmed by the fact that knocking out the Gcv system lead to no growth on glycine only, nor on methanol combined with glycine. The same was observed for the glycine-hydroxymethyl transferase (Shm), which is also essential for the serine cycle and the oxygen tolerant reductive glycine pathway. These results suggest that *K. phaffii* could grow via the oxygen tolerant reductive glycine pathway, if sufficient glycine was produced from methanol or formate and CO_2_, however this is not the case. As the GcvKO strain cannot grow on the combination of methanol and glycine, the limitation must be in the native formation of methylene-THF. Furthermore, evidence was found that not only the glycine cleavage system is limiting, as the overexpression of the Gcv genes did not lead to growth on the combination of methanol and 5% CO_2_. To exclude the fact that growth via the native oxygen tolerant reductive glycine pathway is only inhibited on a transcriptional level, we overexpressed the full pathway from formate to pyruvate with strong methanol inducible promotors and again no growth within 46 days of cultivation could be achieved (Fig. 3 and Supplementary information Fig. S6). As growth on methanol with glycine was slower than on glycine only, we tested the hypothesis that methanol poisoning inhibits growth via alternative methanol assimilation pathways due to *in vivo* formaldehyde accumulation. For that purpose, we cultivated a strain that was additionally *AOX1* deleted to reduce intracellular formaldehyde formation on methanol when combined with CO_2_. As when using formate as carbon source where no formaldehyde is produced at all, no growth was observed in the *AOX1* knockout. This indicates that cell poisoning cannot be the main cause of growth inhibition. In conclusion, the flux limitation for native growth via the proposed pathway has to be either native enzymatic activity, co-factor limitation, or post transcriptional regulatory limitation.

### Further knockout and overexpression strains confirmed the presence of an active reductive glycine pathway in *K. phaffii*

To explore if the oxygen tolerant reductive glycine pathway is the sole alternative methanol and formate assimilation pathway, further targeted gene knockouts were conducted and tested in forward labelling experiments. The enzymatic formate fixation pathway to methylene-THF was deleted by knocking out the *MIS* genes (MisKO strain). Formate and methanol labelling experiments with this strain lead to the following result: after 24 h the fraction of the unlabelled isotopologue M+0 of methionine and serine was only slightly reduced, indicating only minor ^13^C-label incorporation, which was expected with the underlying hypothesis (Fig. 4 a&b). In comparison, the Mis active DasKO strains were heavily labelled at the same timepoint. The decarboxylated (DC) mass spectrometric fragments of these amino acids, which contain only carbon atoms deriving from methanol or formate in the proposed pathway, did not show any increase in ^13^C content. This implies that no other formate or methanol incorporation pathway is active. As the unfragmented amino acids are slightly ^13^C labelled, the label is located in the carboxy groups of the amino acids, which derives from the active carboxylation of the glycine cleavage system (Gcv). Due to the fact that the dissimilation pathway to CO_2_ is still active, ^13^C-CO_2_ is produced intracellularly and can be re-fixed in the Gcv system. Consequently, the labels detected in the unfragmented amino acids come from re-fixed ^13^C-CO_2_, not directly from formate or methanol fixation. To remove doubts as to whether the labelling patterns derived from the experiments with the DasKO strain and the wildtype (described in the prior sections) stem from intracellular ^13^C-CO_2_, we also evaluated the decarboxylated amino acid fragments of these experiments (Fig. 4 and Supplementary information Fig. S7). As they showed ^13^C-labelling, the carbon must indeed come directly from formate. The spontaneous *in vivo* condensation of THF and formaldehyde to methylene-THF was not observed while cultivating the MisKO strain on methanol, hence the enzymatic fixation route via formate is the only alternative methanol assimilation pathway. When knocking out the *SHM* genes, the direct downstream metabolite serine is unlabelled, while upstream methionine and glycine are still labelled (Fig. 4c), which is in accordance with the underlying hypothesis and proves that labels in glycine are not just coming from serine via Shm enzymes.

**Figure 4:**
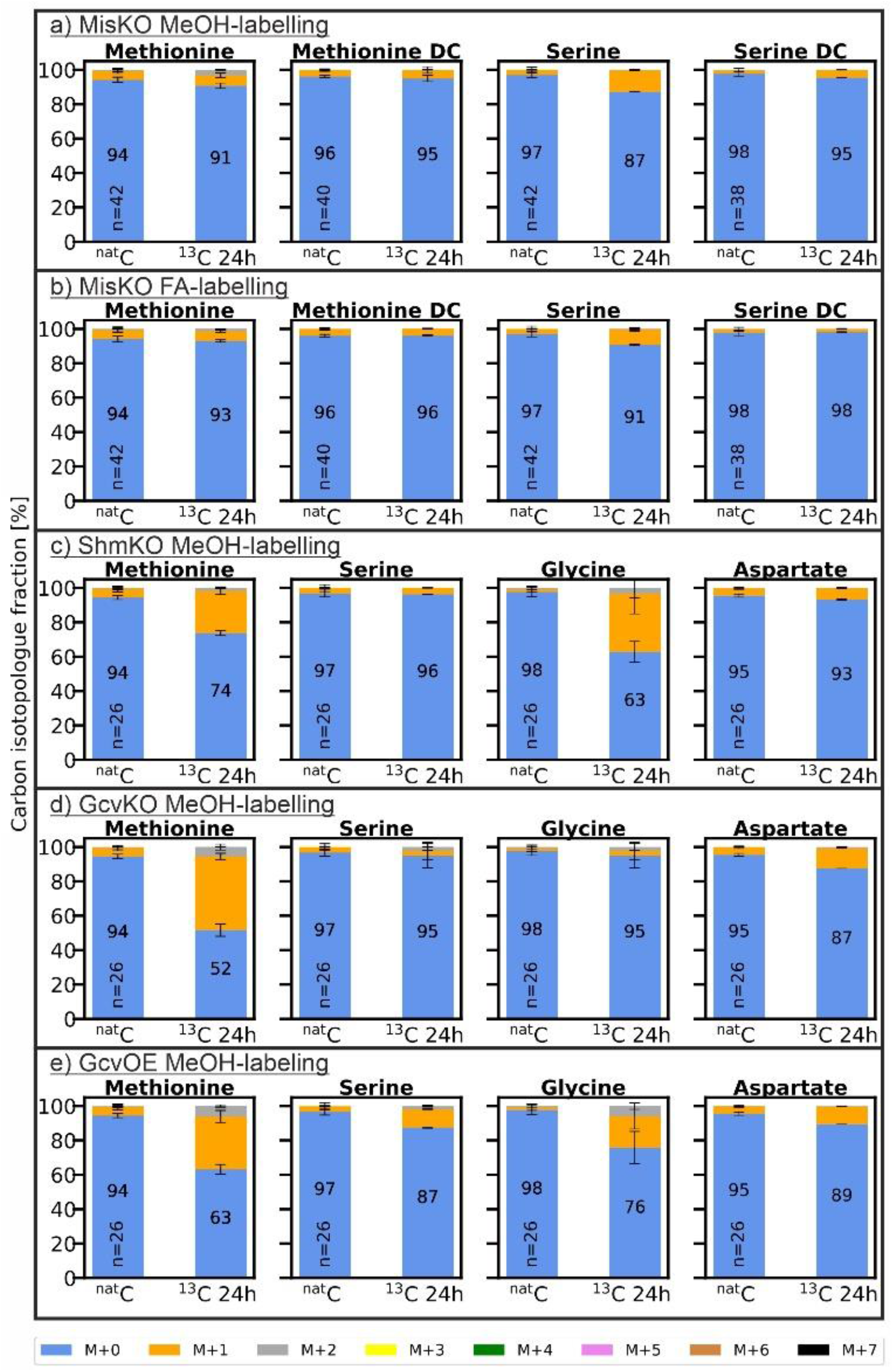
Carbon isotopologue distribution analysis of further knockout strains. As in Figure 2, *K. phaffii* strains (see Table 1) are labelled with different carbon sources (n=2 biological replicates for labelled strains, number of replicates (n) of the ^nat^C controls is indicated in the bar). “BB” in the metabolite name stands for the amino acid backbone, i.e. C1 and C2 only, “DC” stands for the decarboxylated amino acid, i.e. all carbon atoms except C1. a) *das1Δdas2 Δmis1-1Δmis1-2&*3*Δ* strain labelled with ^13^C-methanol for 24h; b) *das1Δdas2 Δmis1-1Δmis1-2&3Δ* strain labelled with ^13^C-sodium formate for 24 h; c) *das1Δdas2Δ shm1Δshm2Δ* strain labelled with ^13^C-methanol for 24 h; d) *das1Δdas2Δ gcv1Δgcv2Δ* strain labelled with ^13^C-methanol for 24 h; e) *das1Δdas2Δ* P_strong_*GCV1&2&3&LPD1* strain labelled with ^13^C-methanol for 24 h. Labelling data of more metabolites see Supplementary information Figure S7 g-l

As in all experiments glycine is less and later labelled than serine, one might suggest that glycine is downstream of serine, raising doubts if glycine is indeed upstream of serine with labels coming from the glycine cleavage (gcv) system into the detected pathway. To underline the involvement of the gcv system and that glycine is upstream of serine, we both deleted and overexpressed the gcv system. In the GcvKO strain, no significant incorporation of ^13^C into glycine could be observed (Fig. 4 d), leading to the conclusion that labels in glycine are only coming via the glycine cleavage system. As serine is downstream in the proposed pathway and can incorporate labels of only one of the two reactions, the ^13^C amount is also less than in the DasKO strain. When overexpressing gcv we could show increased incorporation of ^13^C, more precisely, a reduction of the unlabelled isotopologue M+0 from 90% to 75% after 24 h (Fig. 4 e). Labelling data of further metabolites are found in Supplementary information Fig S7 g-l.

## Discussion

Carbon metabolism enzymes are generally among the most abundant cellular proteins so that they enable high flux rates. Therefore, on any given substrate the most abundant catabolic pathway typically dominates, and less active pathways are easily overlooked in biochemical analyses. In *K. phaffii*, the canonical methanol assimilation pathway is initiated by alcohol oxidase (Aox) to formaldehyde, entering the XuMP cycle to glyceraldehyde 3-phosphate. We have recently described that Aox is not the only enzyme catalysing the first reaction step in the native methanol metabolism. The cytosolic alcohol dehydrogenase (Adh2), known for both synthesis and consumption of ethanol, is also oxidizing methanol to formaldehyde^17^. This reaction is energy conserving by the reduction of NAD^+^ to NADH, but seems to have limitations, as growth was not observable with Adh only. Here, we demonstrate that *K. phaffii* harbours a complete methanol or formate and CO_2_ co-assimilation pathway leading to pyruvate and further to oxaloacetate: the oxygen tolerant reductive glycine pathway, providing precursors for all metabolic routes in yeasts (see Fig. 5).

**Figure 5:**
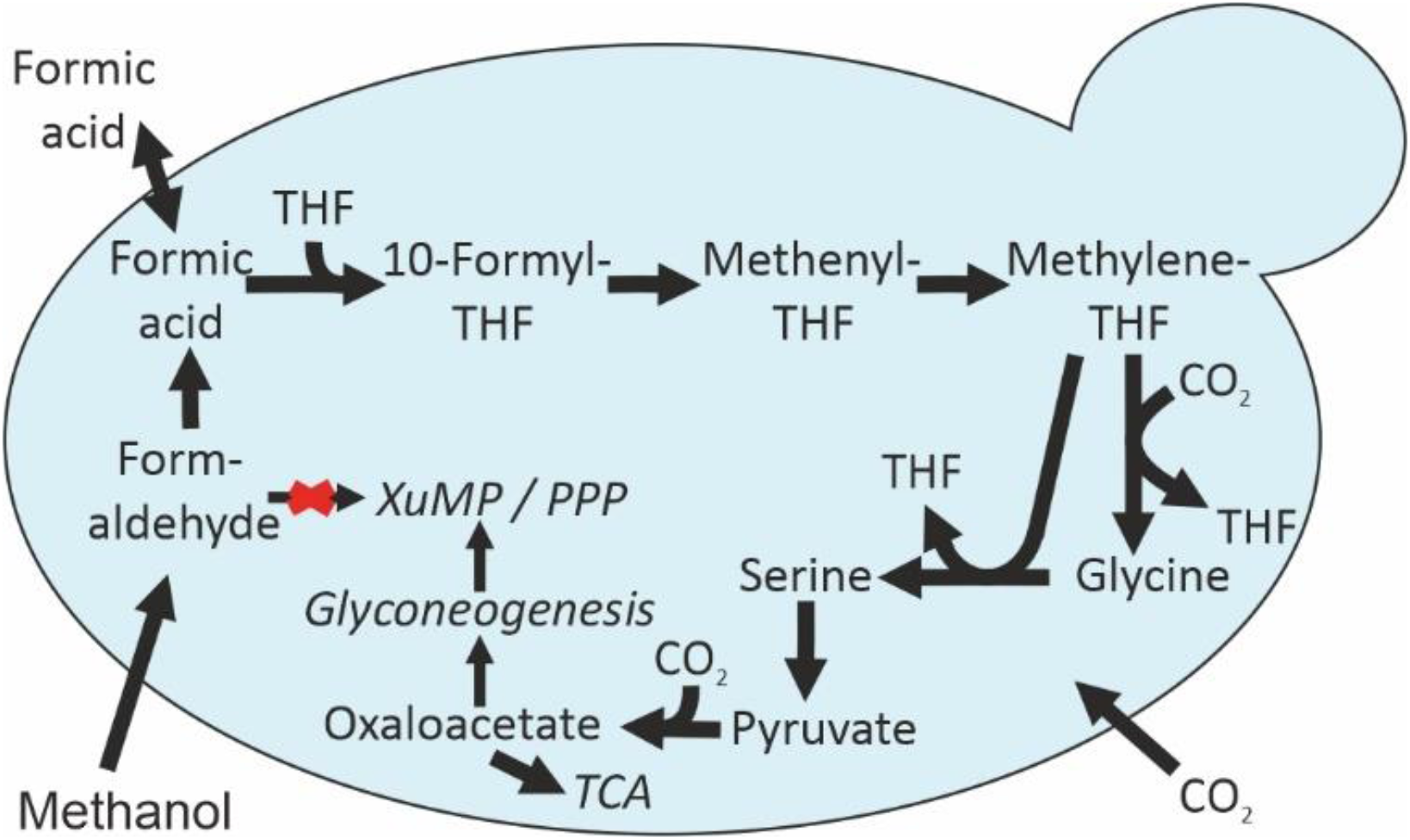
The oxygen tolerant reductive glycine pathway. The oxygen tolerant reductive glycine pathway is the sole natively active alternative methanol assimilation pathway in the yeast *K. phaffii*, being also the sole native formate and a native CO_2_ assimilation pathway. It starts with methanol or formate assimilation and continues via methylene-THF, glycine and serine towards formation of pyruvate and even further towards formation of oxaloacetate. Abbreviations: THF – tetrahydrofolate; TCA - tricarboxylic acid cycle; XuMP – xylulose 5-phosphate pathway; PPP – pentose phosphate pathway

Even if the native metabolic activity of the oxygen tolerant reductive glycine pathway from formate or methanol to pyruvate or oxaloacetate could not be found in nature, it is not entirely new to scientists. The pathway was reported on a metagenomic level for the anaerobic bacterium *Candidatus Phositivorax anaerolimi Phox-21*^31^. However, as this organism is not isolated yet and knowledge on metabolic pathway activity is not accessible, the question remains unsolved if this organism follows the oxygen-sensitive route, just as *D. desulfuricans*, which is also an anaerobic bacterium^29^. In the model organism *E. coli* this oxygen tolerant pathway was integrated and overexpressed as a synthetic route using native genes of other organisms^32–34^. The authors achieved growth on formate, methanol and CO_2_, which proves that this pathway is principally supporting sufficient flux for cell proliferation. When the reductive glycine pathway from formate to glycine is overexpressed in *S. cerevisiae* while blocking all other glycine synthesis routes, the resulting strain is capable of producing sufficient glycine out of formate and CO_2_ to grow on glucose^30^. This concurs with our findings that the glycine cleavage (Gcv) system is reversible and that native Gcv and Mis enzymes show activity in yeasts. The metabolic activity of pyruvate carboxylase in *K. phaffii* is obvious and was metabolically verified by reverse labelling of fully ^13^C-labelled biomass with ^nat^C CO_2_^15^. Our DasKO strain confirmed these findings. In addition, we demonstrated that the glycine cleavage system provides a second natively active CO_2_-fixation reaction.

*K. phaffii* evolved in the direction of using the formaldehyde fixing xylulose 5-phophate pathway as the main methanol utilization pathway, although it has the metabolic and genomic capability to assimilate methanol via the formate-fixing oxygen tolerant reductive glycine pathway. All experiments described in literature, in which growth was achieved via the Gcv system ^30,32,34^ showed that elevated CO_2_ concentrations are a prerequisite. However, as *K. phaffii* was isolated from trees^40^, it evolved under atmospheric CO_2_ concentrations, which seem to be too low to activate glycine synthesis via the Gcv system. The observations of Cruz et al.^30^, who investigated the above-mentioned *S. cerevisiae* strain containing the synthetic route, further support this hypothesis. Because the flux from formate and CO_2_ to glycine was so low, only sufficient glycine to compensate the glycine-autotrophy was produced for growth on glucose. This indicates that flux to glycine for growth solely on formate and CO_2_ without glucose could not be achieved even with the strongly overexpressed pathway. Either the native enzyme activities of yeast are not sufficient and/or the limitation is related to NADPH availability. NADPH is needed in the Mis reaction and produced under glucose feed but not with a feed containing formate only. Also *K. phaffii* produces NADH, not NADPH via the methanol dissimilation pathway ^41^. Our results showed that formaldehyde toxification in the DasKO strain under methanol feed is unlikely to be the only reason for growth inhibition via the detected alternative pathway. This is supported by the findings of Berrios et al.^42^, who described the methanol dissimilation pathway as main formaldehyde detoxification route, which is fully active in all strains used within this study. Both causes seem plausible, as (1) overexpression of NADPH producing reductases increases growth in the above-mentioned *E*.*coli* strains^34^ and (2) growth via the oxygen tolerant reductive glycine pathway could only be achieved in model organisms when using the Mis enzymes of *M. extroquens*^32,34^, which are responsible for the production of methylene-THF. These enzymes support growth in *M. extorquens* via the serine cycle^24,25^ and probably increased their activity in the course of evolution to support faster growth of *M. extorquens* on methanol.

Methanol and formate are considered as valuable substrates for sustainable biotechnology-based production of chemicals as well as food and feed proteins. Both methanol and formate can be produced electrochemically from CO_2_^2,3^. Co-assimilation of CO_2_ with methanol or formate would even allow for direct carbon capturing to reduce greenhouse gas emissions. The discovery of a native functional methanol, formate and CO_2_ assimilation pathway in *K. phaffii* bears huge potential for the design of a chassis cell converting C_1_ substrates to pyruvate and oxaloacetate, two central metabolites from which all carbon backbones of metabolites and microbial biomass can be built. Co-assimilation of CO_2_ with a reduced substrate like methanol is of special interest for the production of rather oxidized molecules like organic acids as it provides the reducing equivalents for CO_2_ assimilation directly in the synthesis pathway^43^.

## Methods

### Plasmid construction

All plasmids were constructed by Golden Gate cloning^13^. For all knockouts, single guide RNA plasmids with the Cas9 protein and linear homologous DNA sequences were constructed for complete gene deletions as described before^14^. Only for the *MIS1-2&3* knockout, a splitmarker cassette was constructed^44^. For the gene deletions of *DAS1* and *DAS2*, the guide RNA plasmids and the homology regions were taken from Gassler et al. 2020^15^. For the knockouts of *GCV1, GCV2, SHM1, SHM2, MIS1-1*, and *MIS1-2&3*, the homology regions were amplified by PCR (NEB, Q5 high-fidelity DNA polymerase) from wildtype genomic DNA of CBS7435 used before^15^. The ligated homologous regions and the kanMX-splitmarker for *MIS1-2&3* were also amplified for transformation with PCR. The single guide RNA recognition site (sequences Supplementary information Table S1) was generated by overlap extension PCR and cloned into CRISPi plasmids^14^. Promoters, terminators and plasmid backbones used for overexpressions were constructed in previous studies^13^. The coding sequences of *GCV1, GCV2, GCV3, LPD1, MIS1-1, SHM1, SHM2, and CHA1* were amplified from the genomic DNA mentioned above. *ADE3* was amplified from genomic DNA of *S. cerevisiae* (S288C). For the final overexpression plasmid constructs see Supplementary information Table S3.

### Strain construction

All knockouts in this study were constructed by CRISPR/Cas9-based homology-directed recombination^14^. 500 ng guide RNA plasmid and 3-5 μg linear homologous DNA sequence were transformed in *K. phaffii* (*P. pastoris*) CBS7435^45^ as previously described^8^. The DasKO strain was constructed within a single transformation with both guide RNA plasmids and linear homologous DNA sequences. This strain served as a basis for all further strain constructions in this study, and additional knockouts were performed consecutively. The *mis1-2&3* knockout was conducted by combining a splitmarker-method^8^ with CRISPR/Cas9 and supplementing YPD plates with 10 mmol L^-1^ hypoxanthine (further details in Supplementary information Chapter 2.2.1). Screening for the correct gene deletion without gene reintegration was performed by colony PCRs using primer pairs binding outside of the homology regions and inside the gene (primer listed in Supplementary information Table S4). For the creation of overexpression strains, 3 μg of the final overexpression plasmids were linearized with either *SmaI* or *AscI* (New England Biolabs) and transformed consecutively into the DasKO strain as mentioned above. Integration in the correct locus was verified with the primer pair listed in Supplementary information Table S4.

### Labelling experiments & metabolic sampling

10 mL YPD precultures were inoculated with a single colony and shaken at 25°C and 180 rpm overnight. A following 100 mL YNB (Sigma-Aldrich GmbH, with 10 gL^-1^ (NH_4_)_2_SO_4_, 0.1 mol L^-1^ potassium-phosphate buffer, pH=6) batch culture was inoculated to an OD_600_ of 1 (25°C, 180 rpm overnight). For methanol and formate forward labelling, 18 g L^-1 nat^C-glycerol (Carl Roth GmbH) and for reverse labelling, 0.79% (v/v) fully labelled ^13^C-glycerol (99atom% ^13^C, CortecNet) were used as carbon sources in the batch cultures. For ShmKO, the batch culture was also conducted on YPD and extended to 48 h batch time to gain sufficient biomass. The MisKO pre and batch cultures were conducted in YPD with 5 mmol L^-1^ hypoxanthine (Sigma-Aldrich GmbH) and 100 mg L^-1^ nourseothricin. Methanol and CO_2_ reverse labelling cultivations were performed in YNB medium with 1% (v/v) ^13^C-methanol (99atom% ^13^C, CortecNet), formate labelling in YNB medium with 30 mmol L^-1 13^C-sodium formate (99atom% ^13^C, Sigma-Aldrich GmbH). For the labelling cultivations with ^nat^C-glycine, glycine (Merck KGaA) was added at a concentration of 20 mmol L^-1^ in addition to methanol. The biomass of the batch cultures was washed and started at an OD_600_ of 50 (which corresponds to 10 g L^-1 nat^C-cell dry weight) for methanol and CO_2_ reverse labelling, or at an OD_600_ of 25 (which corresponds to 5 g L^-1^ cell dry weight) for formate labelling. For experiments with metabolite sampling points at 2 h, 24 h and 72 h, the starting volume was set to 16 mL. ^13^C-methanol was adjusted to 1% (v/v) after 24 h, 48 h and 58 h. ^13^C-formate concentration was adjusted to 30 mmol L^-1^ after 7 h, 17 h, 22 h, 30 h, 41 h and 52 h (for methanol and formate feeding profiles see Supplementary information Figure S8, at each sample point HPLC measurements^17^ were conducted to assess the carbon source concentration). Each metabolic sample had 3 mL, additional samples for methanol measurements 1 mL and for formate measurements 300 μL. Experiments with metabolic sampling at 24 h only had a starting volume of 10 mL. Methanol and formate labelling was conducted at 25 °C at atmospheric gas conditions, CO_2_ reverse labelling at 30°C in 5% ^nat^C-CO_2_ shaken at 180 rpm. All forward labelling experiments were conducted in biological duplicates with parallel ^nat^C-carbon source control experiments.

Metabolic sampling was performed as described before with minor deviations^17,46^. In brief, 1 or 2 mL culture (corresponding to 10 mg ^nat^C-cell dry weight) was quenched in the 4-fold volume of quenching solution (60% methanol (Sigma-Aldrich GmbH), 125 mmol L^-1^ TRIS-HCl (Carl Roth GmbH), 55 mmol L^-1^ NaCl (Merck KGaA, pH 8.2; T = −27 °C). The mixture was immediately vortexed for 4 seconds and rapidly filtered through cellulose acetate filters (0.45 μm, Sartorius Stedim Biotech GmbH). The biomass was washed on the filter with 10 mL of 60% methanol and stored at -70 °C until metabolite extraction.

The labelling experiment of the Mut-strain including sampling and extraction was carried out within the study of Zavec et al. 2021^17^.

### Sample preparation & GC-TOFMS analysis of intracellular metabolites

For gas chromatography – time-of-flight mass spectrometry (GC-TOFMS) isotopologue distribution analysis, the quenched cells were extracted using boiling ethanol extraction as established in our lab for *K. phaffii*^16,47^. Briefly, 4 mL of 75% ethanol (HPLC-grade, Sigma-Aldrich GmbH) at 85°C was added to the cell pellet on the filter. Samples were vortexed, heated for 3 minutes at 85°C and rapidly cooled on dry ice before centrifugation at 4000 g and -20 °C for 10 minutes. The supernatants were dried in a vacuum centrifuge, and dry extracts were stored until analysis at -80°C. Before analysis, the extracts were reconstituted in 1 mL MS-grade water for 30 min at room temperature.

All metabolites were derivatized making use of automated just-in-time online derivatization on a sample preparation robot (MPS2, Gerstel) and analysed via gas chromatography – time-of-flight mass spectrometry (GC-QTOFMS, 7200B, Agilent Technologies).

For some, mainly the phosphorylated metabolites (see Supplementary information Table S8), a method based on ethoximation and trimethylsilylation followed by gas chromatography – chemical-ionization-time-of-flight mass spectrometry (GC-CITOFMS) was used, which was originally designed and developed for the *K. phaffii* wildtype on glucose by Mairinger et al.^36^. Minor adaptions at the GC-MS data acquisition level were made and significant changes were necessary for data analysis. As the cultivation of the XuMP knockout strain (DasKO) on methanol as well as the use of formate as carbon source led to profound changes in analyte concentrations and matrix composition of the labelling samples, the chromatographic and mass spectrometry-related background differed significantly from prior *K. phaffii* samples analysed. Consequently, the method had to be further developed and optimized. Technical changes included (1) slowing down the GC-temperature program in order to reduce interferences (70 °C hold for 1 min, 15 °C min^-1^ to 190 °C, 5 °C min^-1^ to 225 °C, 3 °C min^-1^ to 260 °C, 20 °C min^-1^ to 310 °C, hold for 3 min), and (2) the use of an Agilent split/splitless injector with a splitless gooseneck liner with glass wool (Agilent). 350 μL of the reconstituted extract was dried in 400 μL inserts in 1.5 mL chromatography vials after the addition of 40 μL ethoxyamine hydrochloride in pyridine (*c* = 20 g L^-1^), and vials were crimped with magnetic caps before placing the samples on the autosampler at 7 °C.

In order to cover all metabolites of importance and to extend the biological information by positional information due to specific fragmentation in the electron ionization (EI) ion source, an additional GC-MS based method was implemented, further developed and optimized with regard to measurement and data evaluation. For amino acid and organic acid analysis, TBDMS- (*tert*-butyldimethylsilyl-) derivatization followed by gas chromatography – electron-ionization-time-of-flight mass spectrometry (GC-EI-TOFMS) in split mode (split 1:50, Agilent straight split liners with glass wool) was used^15^. For the analysis of metabolites of low concentration (e.g. glycerate), some derivatized samples were reinjected in splitless mode after changing to an Agilent splitless gooseneck liner with glass wool (see Supplementary information Table S5). This method was used in the study of Gassler et al. 2020^15^ for amino acid analysis only, was extended to cover more metabolites and was applied with minor technical adaptions. More specifically, the final hold time of the temperature programme was extended from 4 to 9 minutes to enable the elution of citrate. 150 μL of the reconstituted extract was dried in inserts for this procedure.

### GC-TOF-MS data evaluation

For isotopologue distribution analysis, the exact masses of a number of adducts and fragments of all target analytes were calculated and used for data analysis in both profile and centroid mode (Supplementary information Table S6&7). Extracted ion chromatograms (EICs) were integrated with Agilent Technologies MassHunter Workstation Quantitative Analysis for TOF (Build 10.1.733.0). The chromatographic traces (EICs) of all m/z ratios, the integration of the peaks and the corresponding mass spectra were visually inspected and rejected if mass spectral or chromatographic interferences were detected, or if peaks were saturated. In case the automated integration was not satisfactory, peaks were reintegrated manually. The resulting peak areas were corrected for natural heavy isotopes of H, N, O, Si, S atoms of the derivatized molecule as well as C isotopes coming from derivatization (non-metabolite carbon atoms) using the ICT correction toolbox v.0.04 ^48^. Carbon isotopologue fractions were calculated (with *n* = number of carbon atoms in the metabolite, *A*_*i*_ = ICT corrected peak area of isotopologue i, i.e. an isotopologue containing i numbers of ^13^C atoms) according to equation (1):

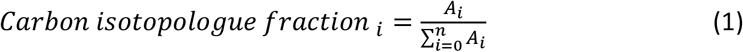

Several fragments and adducts were evaluated in profile and centroid mode for each metabolite, and the corresponding ^13^C carbon isotopologue fractions were calculated. The decision on which fragments or adducts to finally evaluate for different time points and strains was based on the trueness of the results. For this purpose, the isotopologue fractions obtained for ^nat^C labelled metabolites measured in ^nat^C extracts (average of 2 biological replicates) were compared with the calculated natural isotopologue fractions (calculated via https://www.envipat.eawag.ch/ ^49^). This was done in 2 ways: (1) the uncorrected measured ^nat^C isotopologue distributions (not ICT corrected, hence corresponding to the isotopologue fraction of the whole derivatized molecule with *A*_*i*_ being the uncorrected peak areas of respective isotopologues) were compared with the isotopologue fractions calculated via Envipat for the whole, derivatized compound and (2) the carbon isotopologue fractions as obtained after ICT correction were compared to the isotopologue fractions calculated via Envipat for the respective number of carbon atoms in the metabolite. Fragments or adducts were only considered as fit for further consideration if both the deviation between measured and calculated isotopologue fraction and the deviation between measured and calculated carbon isotopologue fraction was less than 5%. For each strain, time point and metabolite the fragment or adduct in centroid or profile mode was selected for further data interpretation that had the lowest deviations of the (carbon) isotopologue fractions from the calculated theoretical values and the lowest standard deviation for ^nat^C and ^13^C replicates (chosen adducts/fragments and type of data (profile/centroid) are summarized in Supplementary information Table S8). ^13^C carbon isotopologue fraction data in Fig. 2 and Fig. 4 are shown as the average of biological duplicates with error bars representing the standard deviation calculated from duplicates multiplied by the correction factor of 1.253314, as proposed by Wegscheider et al.^50^. ^nat^C carbon isotopologue fraction data are presented as mean and standard deviation of n replicates of the selected fragments or adducts from naturally distributed samples.

### Genome mining for alternative methanol assimilation genes

Genes and enzymes for the reactions of interest were either found in the database KEGG (https://www.genome.jp/kegg/pathway.html#metabolism) or alternatively enzymes were searched via the corresponding metabolites in Expasy’s chemical compound search (https://enzyme.expasy.org/enzyme-bycompound.html); related genes were found via the Pichia genome database^45^ (www.pichiagenome.org). Non-annotated enzymes were further checked for their presence in *S. cerevisiae* (https://www.yeastgenome.org/). The homologous gene was either directly found in the Pichia genome database or identified by NCBI’s BLAST, as shown in Supplementary information chapter 1.1.1) (https://blast.ncbi.nlm.nih.gov/Blast.cgi?PAGE_TYPE=BlastSearch).

### Growth experiments

Cultivations were conducted in 100 or 250 mL shake flasks with working volumes between 8 and 25 mL, containing YNB medium. Precultures and batch cultures of the knockout strains were conducted as in the labelling experiments above. For the overexpression strains the batch culture was skipped. The carbon sources of the test culture were 20 mmol L^-1^ glycine, 20 mmol L^-1^ serine (Carl Roth GmbH), 30 mmol L^-1^ sodium formate (Carl Roth GmbH) or 4 g L^-1^ methanol (Carl Roth GmbH) for the first 24 h, concentration was then maintained at 8 g L^-1^, by feeding every 2-3 days. Cultures were started with an optical density at 600 nm (OD_600_) of 0.5 or 1. Flasks were incubated at 25 °C in atmosphere, at 30 °C and 5% CO_2,_and at 25 °C at 10% CO_2_ while shaking with 180 rpm. OD_600_ and methanol or formate concentrations (HPLC measurements^17^) were monitored every 2-3 days. The experiments were conducted in duplicates.

## Acknowledgements

This work was supported by the Austrian Science Fund (FWF W1224, Doctoral Program on Biomolecular Technology of Proteins (BioToP)). EQ-BOKU VIBT GmbH and the BOKU Core Facility Mass Spectrometry are acknowledged for providing mass-spectrometry equipment. We kindly thank P. Tondl for help with GC-MS measurements, H. Rußmayer and L. Lutz for help with labelling experiments, M. Baumschabl for help with data visualization with python, T. Gassler for scientific input and C. Wachtel for the help concerning the ShmKO strain construction.

## Authors contributions

D.M. and S.H. conceived and initiated the project. B.M.M. and D.M. designed the experiments. B.M.M. carried out all experiments, plasmid and strain constructions, measurements, method adaptions & development and all data analysis. C.T. supervised and directed all GC-MS related parts. B.M.M. wrote the original draft and was involved in reviewing and editing the manuscript. All authors reviewed, edited and approved the final manuscript.

