## Supplementary information for "The oxygen tolerant reductive glycine pathway in eukaryotes – a native methanol, formate and CO_2_ assimilation pathway in the yeast *Komagataella phaffii*"

### Overview of Tables

### Overview of Figures

### 1 Supplementary Results

#### 1.1 Bioinformatic analysis

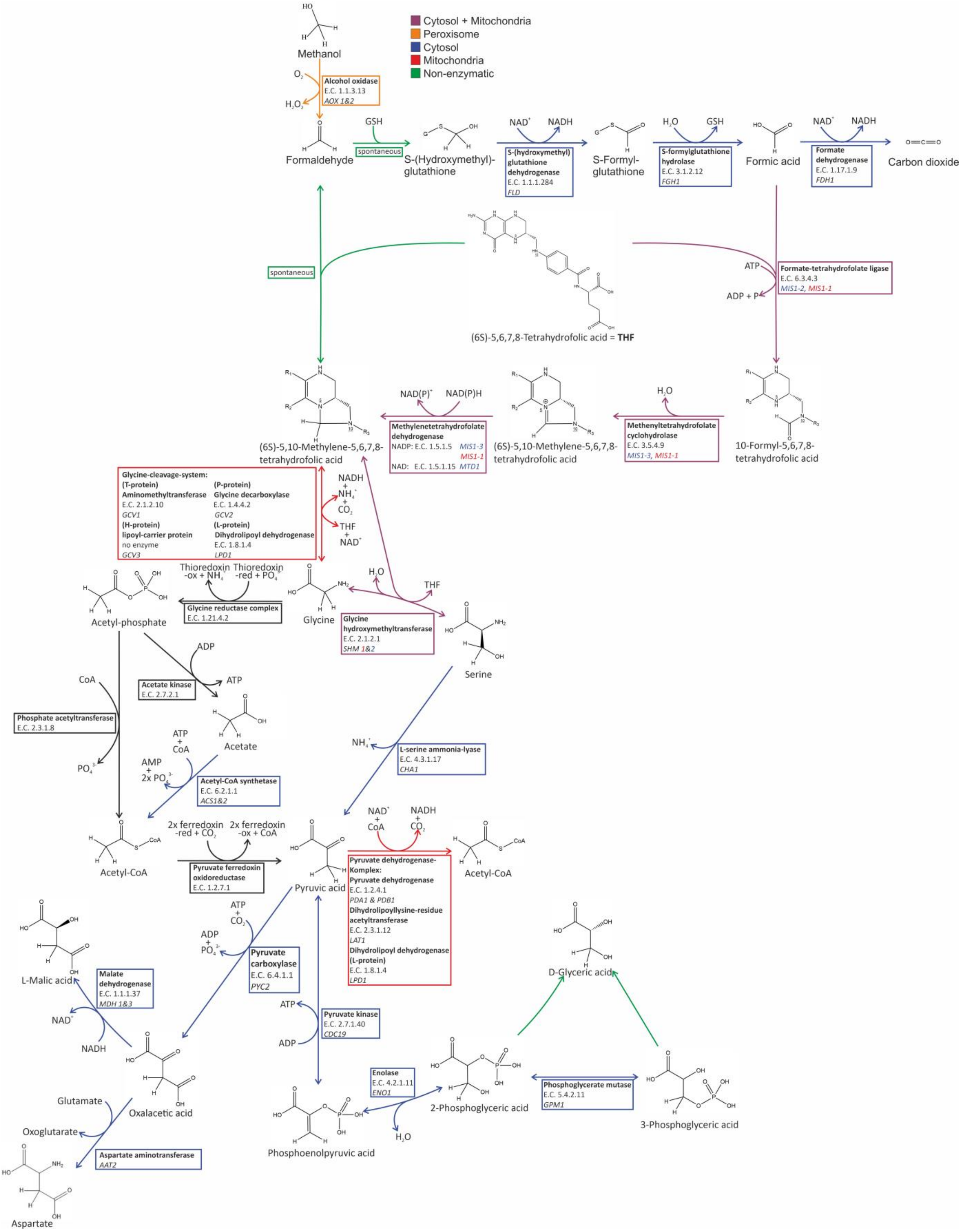

**Figure S1: Detailed reductive glycine pathway** with structures, enzyme annotations, gene annotations and compartment localization in *K. phaffii*; $O_2$-sensitive through acetyl-P, $O_2$- tolerant pathway over serine

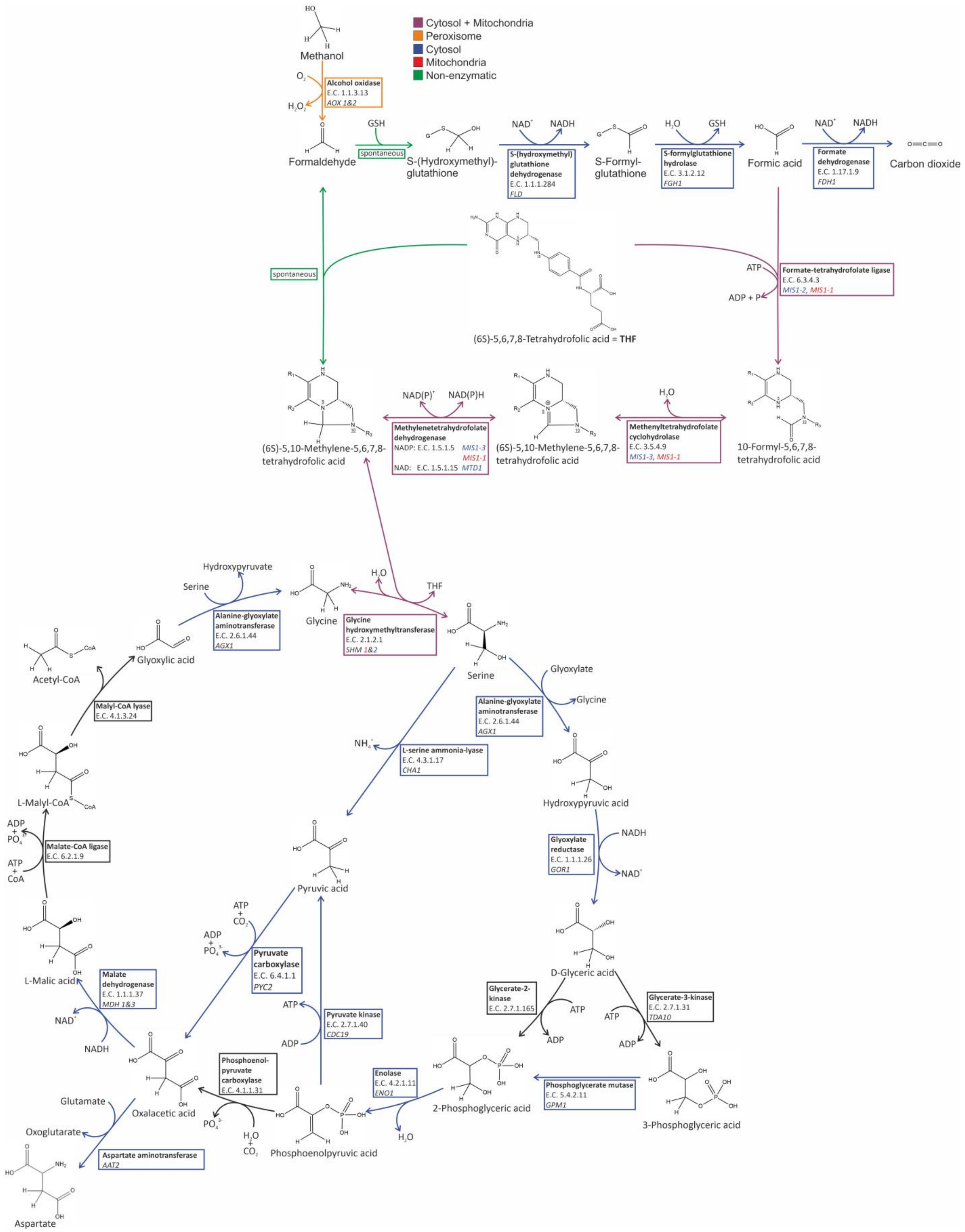

**Figure S2: Detailed serine cycle** with structures, enzyme annotations, gene annotations and compartment localization in *K. phaffii*; the native pathway is shown as a cycle, alternative routes and shortcuts are shown as straight reactions

#### 1.1.1 Cytosolic MIS gene separation

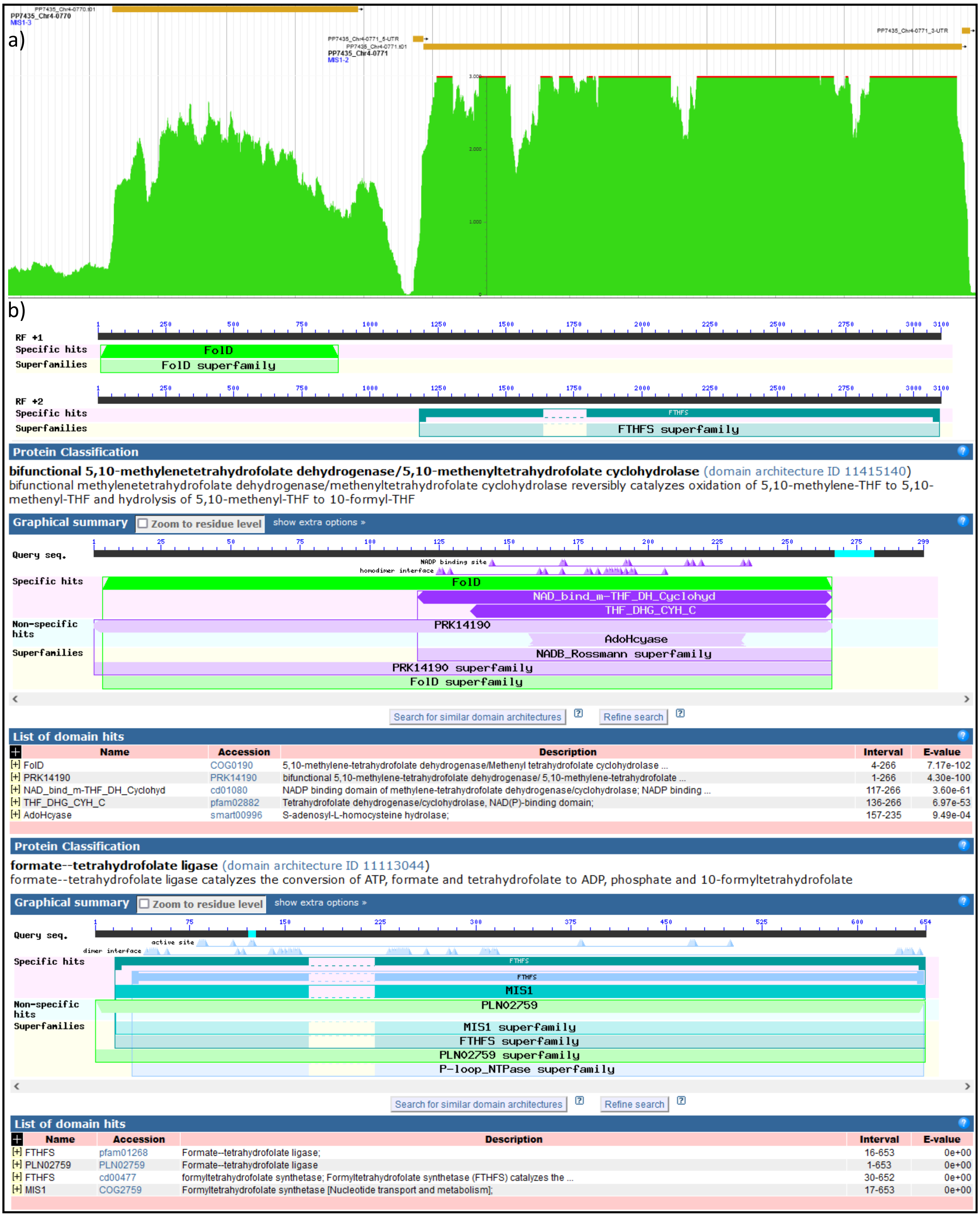

**Figure S3: Cytosolic homologs MIS1-2&3 of K. phaffii are separated** a) mRNA enriched coverage histogram of *MIS1-2&3* of *K.phaffii* (http://pichiagenome-ext.boku.ac.at:8080/apex/f?p=100:23:1507211694349::NO)
b) protein alignment with NCBI´s BLASTp of *MIS1-2* and *MIS1-3* (https://www.ncbi.nlm.nih.gov/Structure/cdd/wrpsb.cgi?RID=B8U6Y2FS013&mode=all, https://www.ncbi.nlm.nih.gov/Structure/cdd/wrpsb.cgi?RID=B8U7N3P2013&mode=all)

The cytosolic MIS gene of *K. phaffii* is split into *MIS1-3* (methylenetetrahydrofolate dehydrogenase & methenyltetrahydrofolate cyclohydrolase) and *MIS1-2* (formate-tetrahydrofolate ligase), as can be seen in

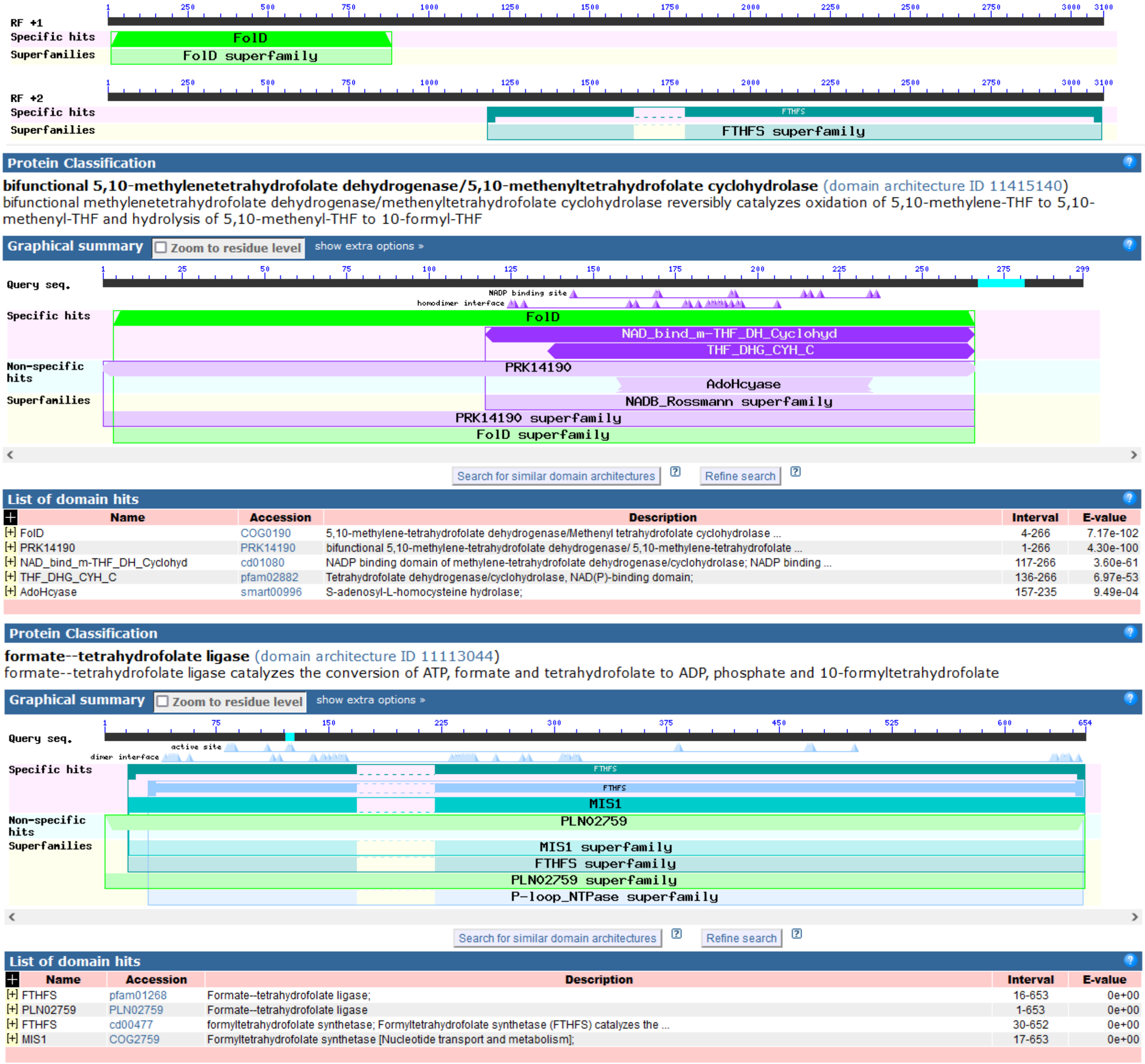

**Figure S3**b. This split results in a higher expression of the second gene *MIS1-2* (Figure S3a). These enzymes are generally essential for native growth (see chapter: 2.2.1), as they are involved in the formation of 10-formyl-tetrahydrofolate, which is necessary for de-novo purine synthesis. Other yeasts, such as *S. cerevisiae* with *ADE3,* still have these enyzmes expressed in one gene (Figure S5). The mitochondrial homolog *MIS1-1* in *K. phaffii* is not split (Figure S4). *MIS1-1* seems to be not intensely involved in de-novo purine synthesis as its knockout does not influence growth on substrates without any hypoxanthine supplementation. Nonetheless, the cytosolic version seems to be of significant importance, as when knocked out growth is not feasible unless hypoxanthine supplementation (see chapter 2.2.1). Higher expression of formate-tetrahydrofolate ligase due to the gene separation might have given *K. phaffii* an evolutionary advantage, as more 10-formyl-tetrahydrofolate might be formed when grown on methanol via formate fixation, which would in turn lead to faster de-novo purine synthesis.

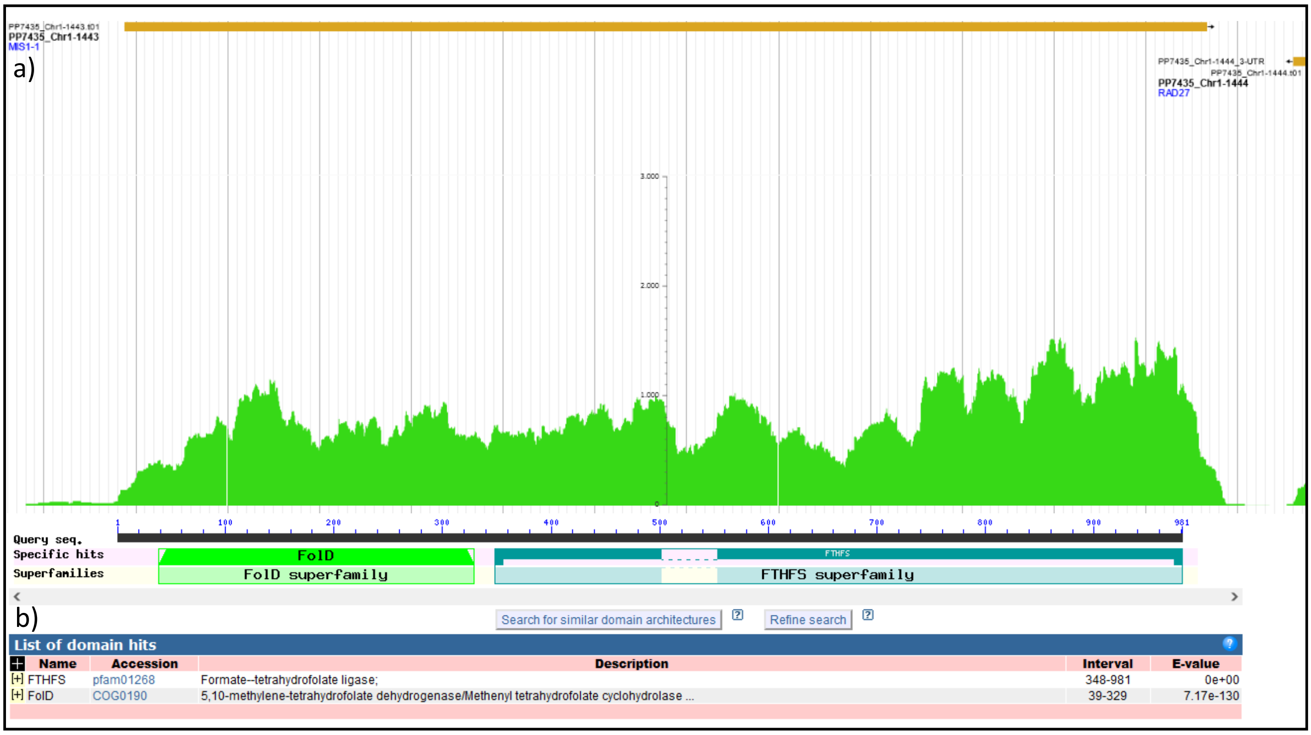

**Figure S4: Mitochondrial homolog *MIS1-1* of *K. phaffii* has no separation**: a) mRNA enriched coverage histogram of *MIS1-1* of *K. phaffii* (http://pichiagenome-ext.boku.ac.at:8080/apex/f?p=100:23:1507211694349::NO) b) protein alignment with NCBI´s BLASTp of MIS1-1 (https://www.ncbi.nlm.nih.gov/Structure/cdd/wrpsb.cgi?RID=B8WME1V6016&mode=all)

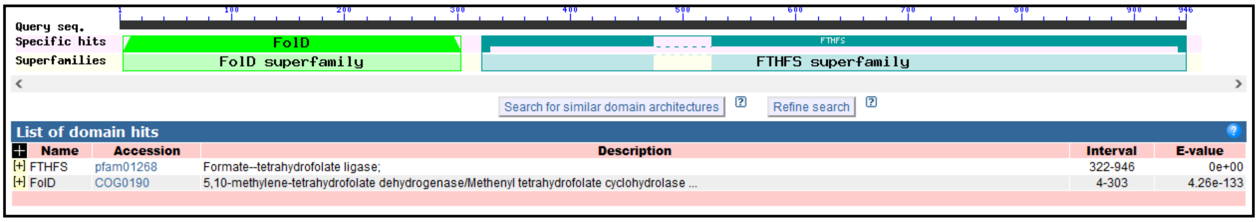

**Figure S5: Cytosolic homolog *ADE3* of *S. cerevisiae* has no separation**: protein alignment with NCBI´s BLASTp of ADE3 (https://www.ncbi.nlm.nih.gov/Structure/cdd/wrpsb.cgi?RID=B8WZ38XT013&mode=all)

#### 1.2 Growth analysis

**Table S1:** Additional strains not mentioned in the main manuscript, their abbreviations and genotypes

| Strain name | Genotype | Source |
|---|---|---|
| Single MisKO | CBS7435 *Δdas1Δdas2 Δmis1-1* | This study |
| mitoOtRedGlyOE | CBS7435 *Δdas1Δdas2* $P_{FDH1}$*GCV1* $P_{DAS2}$*GCV2* $P_{AOX1}$*LPD1* $P_{DAS1}$*GCV3* $P_{DAS1}$*MIS1-1* $P_{AOX1}$*SHM1* $P_{DAS2}$*CHA1* | This study |

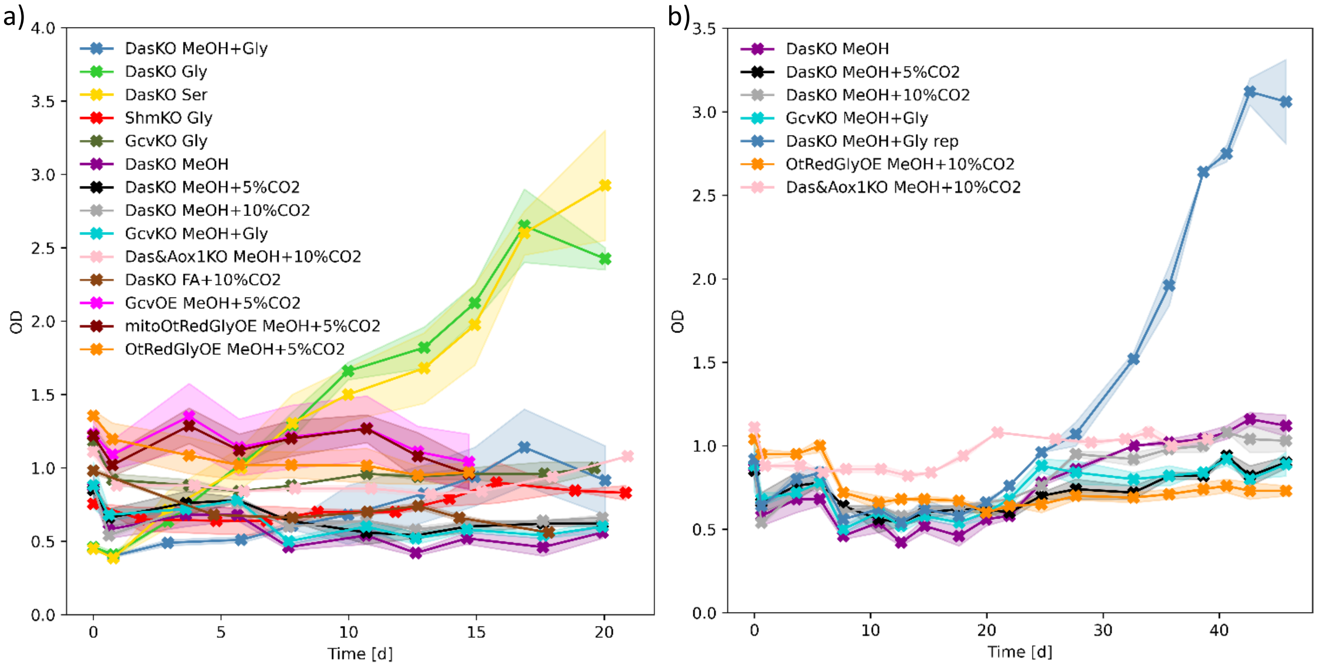

**Figure S6: Growth experiment results:** a) non-normalized $OD_{600}$ of main manuscript, n=2 for knockouts, n= b) long cultivations and repeatability study

#### 1.3 Labelling experiments

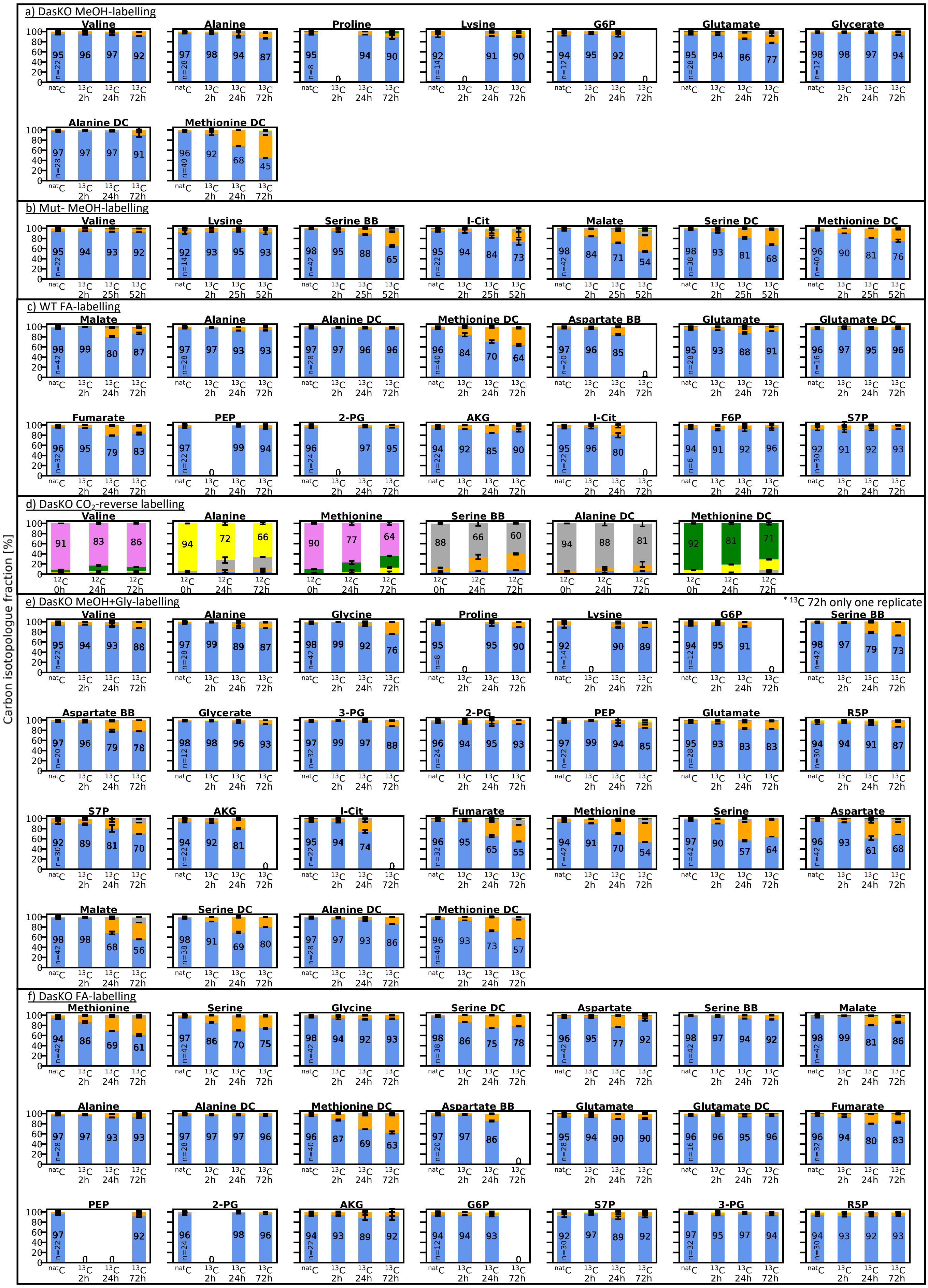

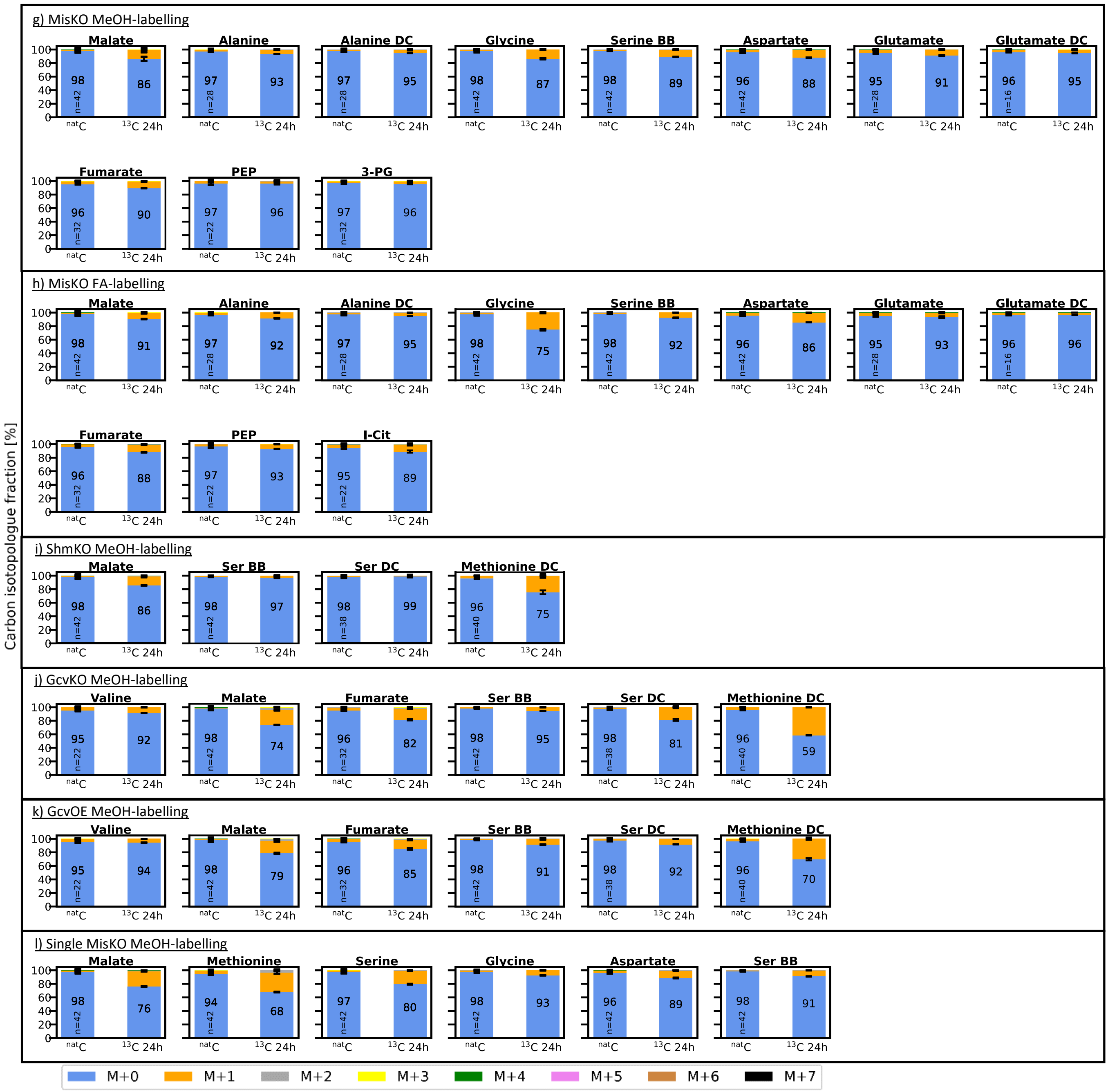

**Figure S7: Additional results of carbon isotopologue distribution analysis** (in addition to the results shown in the main manuscript)

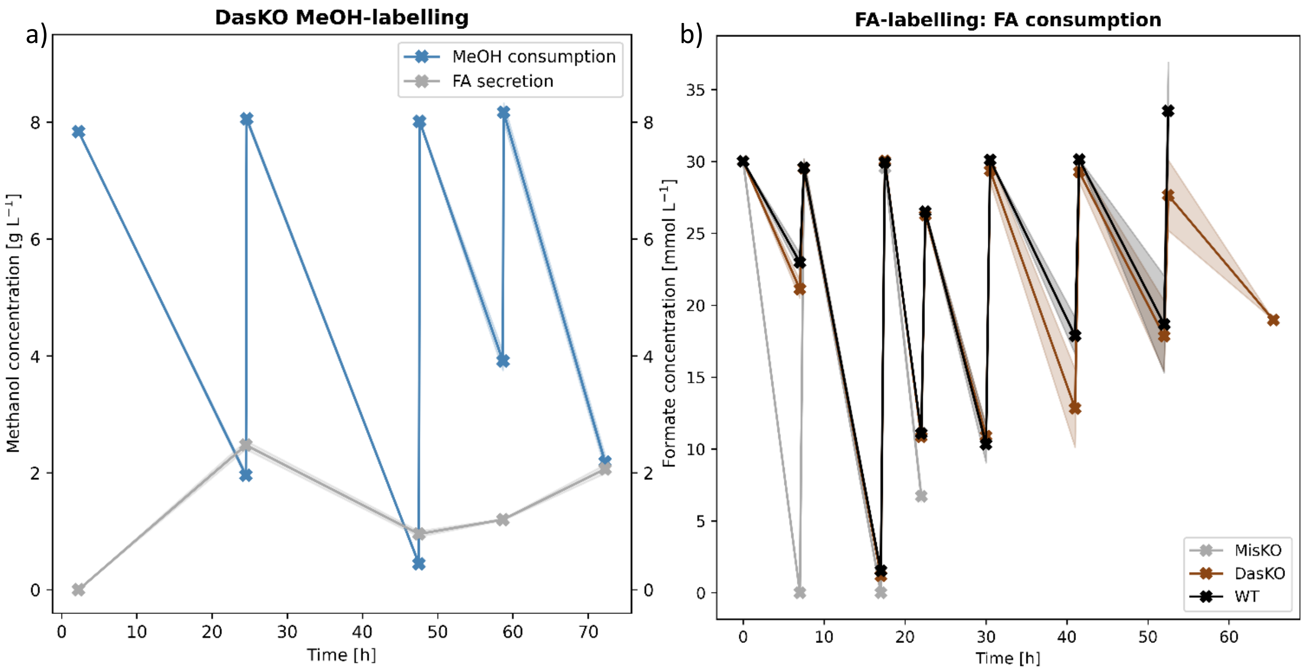

**Figure S8: Carbon source consumption and secretion profile for labelling experiments:** methanol and formate concentration timelines for a) methanol consumption during DasKO methanol-labelling and corresponding formate secretion/ reconsumption in the supernatant, b) formate consumption during formate labelling

##### 1.3.1 Degradation and interconversion of metabolites

Analyte interconversion reactions are an issue in metabolomics, as they falsify quantitative as well as labelling results and mislead their interpretation. As an example, the measured concentration or the labelling pattern of a targeted analyte can either stem from the actual intracellular metabolite under investigation or from the chemical degradation of another component of the cell extract resulting in this metabolite or it can be the sum of both. Interconversion reactions can occur anytime between quenching of the cells and ionization in the mass spectrometer. Especially harsh conditions applying high temperature or non-aqueous and/or non-buffered conditions increase the likelihood of such chemical interconversions. We applied such conditions also in our protocol, especially during boiling ethanol cell extraction and during derivatization and injection for GC-MS. Harsh extraction conditions cannot be avoided as they were shown to be necessary to extract metabolites with high recovery from yeast[1]. Furthermore, GC-MS is still the gold standard for the separation of sugar and sugar phosphate isomers as well as other phosphorylated metabolites and demands for derivatization.

Special attention has to be paid to interconversion reactions when the unphosphorylated or decarboxylated analogue of a metabolite is targeted in quantitative or tracer analysis. As pyruvate and glycerate are of major importance in our study, as branch points for different possible pathways, we focused on these two metabolites. For tracking interconversion, we measured single standards of potential educts. The chromatograms and mass spectra of the respective degradation products are shown in **Error! Reference source not found.** to **Error! Reference source not found.**. The identity of the degradation products was confirmed by comparing them to authentic standards of the respective metabolites.

Glycerate is a metabolite involved in the serine cycle and is analysed for this study via TBDMS GC-EI-TOFMS with splitless injection. Analysis with EtOx/TMS GC-CI-TOFMS is not possible due to interference with other metabolites in the extract. As degradation of 2-PG and 3-PG was suspected due to prior results showing instability of 2-PG even in samples stored at -80°C, we investigated the stability of these metabolites with regard to glycerate formation. Our results confirm, that glycerate is, besides being an important metabolite, a degradation product of 2-phosphoglycerate (2-PG) and, to a lower extent, also of 3-phosphoglycerate (3-PG) as depicted in **Error! Reference source not found.**. It has to be pointed out that although the glycerate concentration is close to LOD in the chromatograms shown in Figure S9, the problem is more severe, as the metabolite glycerate is low in concentration and hence measured with splitless injection (see Table S7), while the chromatograms in Figure 9 were recorded with a 1:50 split injection. Therefore, it needs to be taken into account, that the labelling pattern of glycerate can be falsified by a contribution of the labelling patterns of 2-PG and 3-PG. By comparing the labelling degrees of the detected glycerate peak with the labelling degree of 2-PG and 3-PG, it can nevertheless be evaluated, whether the metabolite glycerate is labelled to a higher extent than 2-PG or 3-PG. If this was the case, the higher labelling degree could only stem from metabolically active pathways, in which glycerate is metabolically upstream of its phosphorylated analogues. In our case this would indicate that the natural serine cycle is active. However, this is not the case in our findings, as glycerate, 2-PG and 3-PG show similar labelling patterns. Hence the labelling pattern of the detected glycerate peak could also stem from 2-PG and 3-PG degradation only.

The problem of interconversion is even more severe if degradation products stem from two different educts, which are involved in separate pathways. This is the case for pyruvate. Pyruvate is the decarboxylation product of oxaloacetate (see **Error! Reference source not found.**a), which is involved in the TCA cycle as well as in the oxygen tolerant reductive glycine pathway and the serine cycle pathway. Pyruvate is also the degradation product of phosphoenolpyruvate (PEP) (see **Error! Reference source not found.**b), which is involved in glycolysis and gluconeogenesis. The pyruvate labelling pattern could therefore show contributions of up to three different labelling patterns: the actual pattern of the intracellular metabolite pyruvate, the pattern of PEP and the pattern of oxaloacetate. As it is impossible to distinguish between these contributions or quantify them, we did not evaluate the labelling pattern of pyruvate.

As can be seen in **Error! Reference source not found.**, dephosphorylation of dihydroxyacetonephosphate (DHAP) to dihydroxyacetone (DHA) and glyceraldehydephosphate (GAP) to glyceraldehyde (GA) was also observed. Therefore, also DHA and GA were not evaluated. DHAP and GAP could not be evaluated as they were either below the limit of detection or the mass spectra showed severe interferences from other constituents of the cell extracts. As a matter of fact, such interconversion reaction could also occur in the case of other metabolites, e.g. dephosphorylation of other phosphorylated metabolites, and deamination of asparagine to aspartate[2,3]. As asparagine is always considered to be metabolically downstream of aspartate, pathway interpretations based on isotopologue distribution analysis are not hampered.

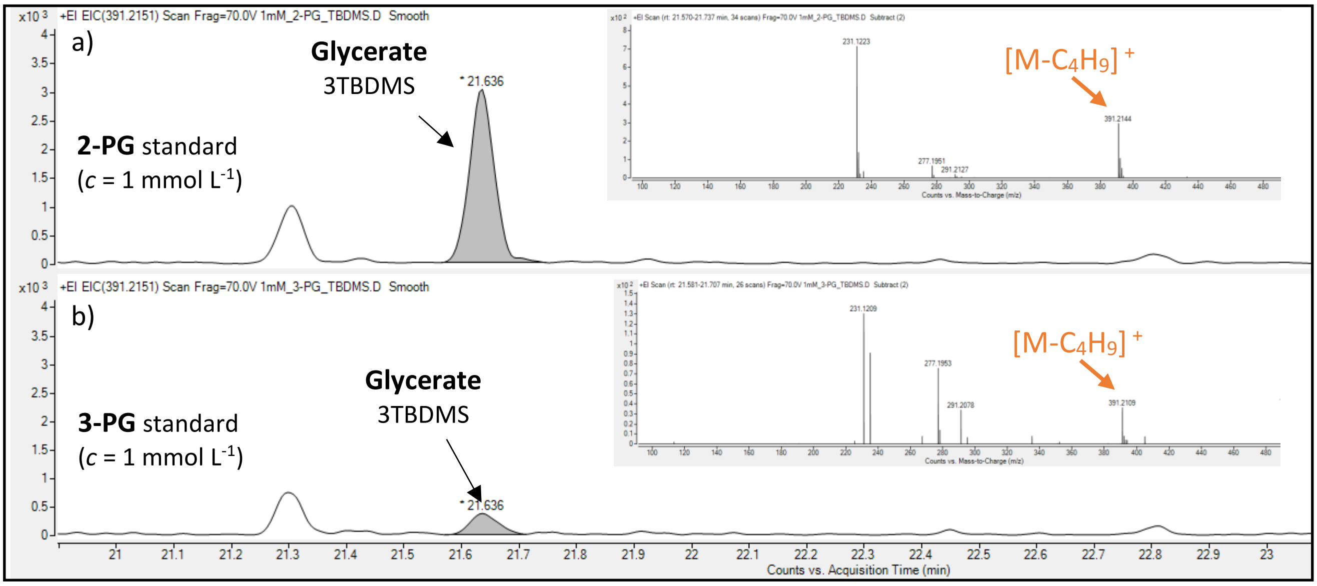

**Figure S9: 2-PG and 3-PG degradation to glycerate shown by TBDMS GC-EI-TOFMS (split injection 1:50)**: chromatogram and mass spectrum of the glycerate peak a) in a 2-PG standard ($c$ = 1 mmol $L^{-1}$) and b) in a 3-PG standard ($c$ = 1 mmol $L^{-1}$)

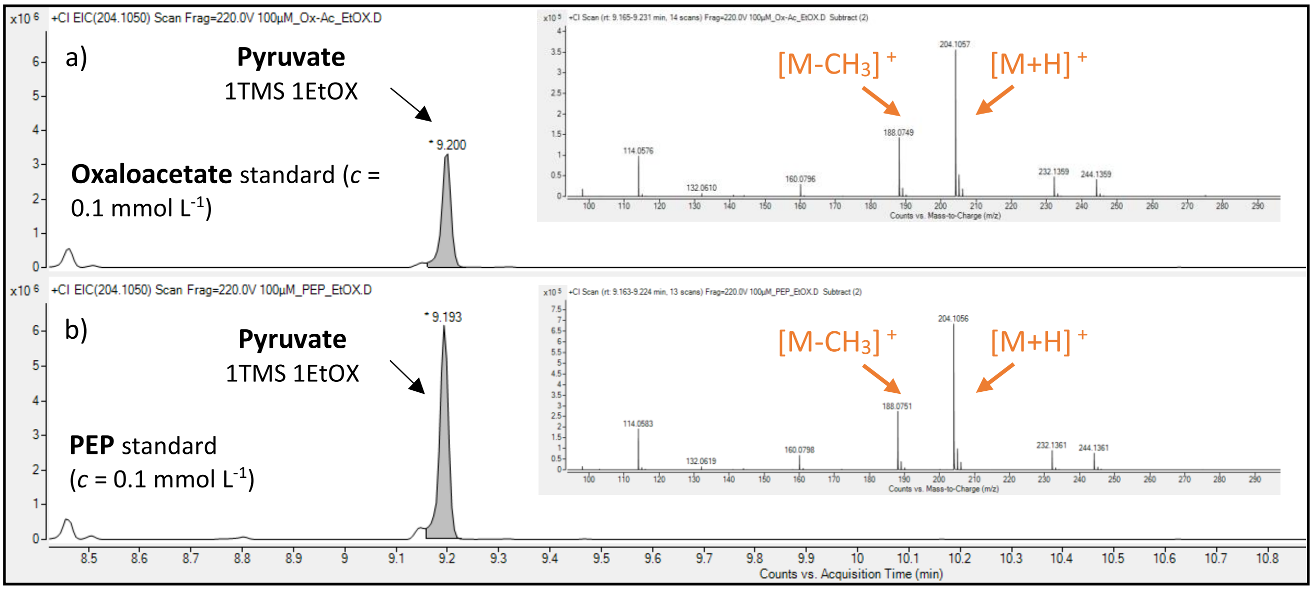

**Figure S10: Oxaloacetate and PEP degradation to pyruvate shown by EtOx/TMS GC-CI-TOFMS (splitless injection)**: chromatogram and mass spectrum of the pyruvate peak a) in an oxaloacetate standard ($c$ = 0.1 mmol $L^{-1}$) and b) in a phosphoenolpyruvate standard ($c$ = 0.1 mmol $L^{-1}$)

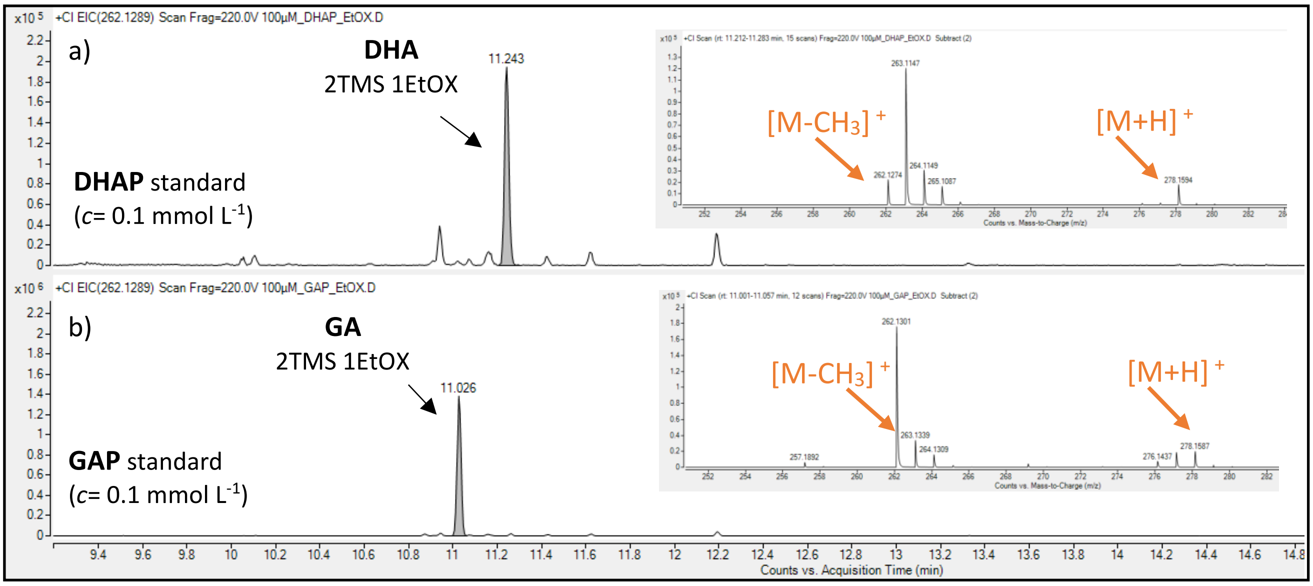

**Figure S11: GAP & DHAP degradation to GA & DHA shown by EtOx/TMS GC-CI-TOFMS (splitless injection)**: chromatogram and mass spectrum a) of the dihydroxyacetone peak in a dihydroxyacetone-phosphate standard ($c$ = 0.1 mmol $L^{-1}$) and b) of the glyceraldehyde peak in a glyceraldehydephosphate standard ($c$ = 0.1 mmol $L^{-1}$)

### 2 Supplementary Methods

#### 2.1 Plasmid construction

**Table S2:** single guide RNA recognition site sequences

| | |
|---|---|
| *GCV1* | TCTTGTAGAAGATACAGCACGGG |
| *GCV2* | GTTGAGATCACACAGAGTGAAGG |
| *SHM1* | CTTCTACATTTCCTGTTCCGGGG |
| *SHM2* | TCAAGAATGAGATTAGCGCCTGG |
| *MIS1-1* | AGCACCGTTAGAAGGCTCGCTGG |
| *MIS1-2&3* | ATGACAAGGGAGAGATTGAAGGG |

**Table S3:** Overexpression Plasmids

| |
|---|
| BB3rN_pFDH1_GCV1m_RPL2Att_pDAS2_GCV2m_RPP1Btt_pAOX_LPD1m_RPS2tt_pDAS1_GCV3m_IDP1tt |
| BB3eH_pDAS1_MIS1-1m_IDP1tt_pAOX1_SHM1_RPS2tt_pDAS2_CHA1_RPP1Btt |
| BB3aK_pDAS1_ADE3_IDP1tt_pAOX1_SHM2_RPS2tt |

#### 2.2 Strain construction

##### 2.2.1 Combining splitmarker cassettes and CRISPR/Cas9 for improved knockout-yields for the lethal *mis1-2&3* gene deletion

The mitochondrial homologous gene *mis1-1*, which encodes for formate-tetrahydrofole ligase, methenyltetrahydrofolate cyclohydrolase and methylenetetrahydrofolate reductase, can be knocked out with the CRISPR/Cas9-based homology directed recombination. The cytosolic *mis1-2&3* version can neither be knocked out with the above mentioned CRISPR system, nor a recombination based splitmarker method, nor the combination of both, even if the mitochondrial homolog is still present. We screened over 600 clones with the CRISPR System, but only 3 knockouts with recombination were found. When applying the splitmarker, 14 knockouts out of 48 clones were detected, but all with reintegrations. If the splitmarker method and the CRISPR-plasmid with Cas9 and guide RNA is transformed together and selected for both, the yield could be increased to 18 knockout clones with reintegrations out of 42 clones. We observed that the *mis1-2&3* gene can only be knocked out by supplementing YPD medium with 10 mmol $L^{-1}$ hypoxanthine, as the *mis* genes are involved in purine de-novo synthesis and a knockout of these is therefore lethal. When supplementing with hypoxanthine, 8 out of 57 clones which were transformed with the splitmarker system only had a knockout, but only one clone without reintegration. When transforming with the the splitmarker system and the CRISPR-plasmid, but selecting for the splitmarker only, the yield was increased to 5 knockouts, with 1 out of 13 screened clones without reintegration. The best yield was achieved by combining splitmarker and CRISPR-plasmid and selecting for both of them (selecting with 2 resistances increased the incubation time of the traffo plate from 2 to 3 days at 30°C). 32 knockouts with 25 clones without reintegration out of 38 clones were detected. These findings show that the combination of the splitmarker method and the CRISPR-plasmid with Cas9 and guide RNA increases the yield of difficult or lethal knockouts. The *mis1-2&3* knockout without reintegration grow slower and cannot grow when hypoxanthine is not supplemented. This knockout was also tested for capability of growth on YNB media with 18g $L^{-1}$ glycerine and 5 mmol $L^{-1}$ hypoxanthine, but growth stopped after one doubling. Additionally the doubling took 24 h, indicating that hypoxanthine is important for the mis1-2&3 knockouts but other ingredients of YPD are necessary for continuous growth.

**Table S4:** Primer list

| | |
|---|---|
| DAS1_Seq_ex_fw | ATTCTGTCGAAAATGGAAGCG |
| DAS1_Seq_ex_rev | CACTTGCATCACTGGCT |
| DAS1_Seq_int_fw | GGTCATCAAAACCTTCCGTT |
| DAS1_Seq_int_rev | AGCCTTGATAGAGTTGACATATTG |
| DAS2_Seq_ex_fw | ATGAAAGGGTTACGGGTGTT |
| DAS2_Seq_ex_rev | TGCTGGCTGGTGTATCTCTC |
| DAS2_Seq_int_fw | CGGTGAATTCGTAAAGGATTTGA |
| DAS2_Seq_int_rev | GCAGTATCGACACAAGATGAC |
| GCV1_Seq_ex_fw | CTTGTATTTTCCTTTCAGGGGG |
| GCV1_Seq_ex_rev | GTTGGGAGATACTCTTAATAGTTTCC |
| GCV1_Seq_int_fw | GAGTAGAAGCTCTTATCAAAACTCC |
| GCV1_Seq_int_rev | GCTACAAAAGGAAGTTTGGC |
| GCV2_Seq_ex_fw | GTCTACAAGATTGGTGGTATCG |
| GCV2_Seq_ex_rev | GAAGGAGTTATGAACCTAAACTGG |
| GCV2_Seq_int_fw | CACAGCAAGTTCATGTCTCC |
| GCV2_Seq_int_rev | GAGATCACACAGAGTGAAGG |
| SHM1_Seq_ex_fw | CATTGTGGAACGGTATTTGC |
| SHM1_Seq_ex_rev | CATTCTAAGCATTGGAAAAGATCG |
| SHM1_Seq_int_fw | GAACAGGAAATGTAGAAGCCC |
| SHM1_Seq_int_rev | CTTAGTCGTAACCTGCAAAGG |
| SHM2_Seq_ex_fw | GTTGATGCTGTTTGTTCAGC |
| SHM2_Seq_ex_rev | CTTATCAAGGTTAACGGTTCACC |
| SHM2_Seq_int_fw | CTAAACAGACAATGGGAATTCTCC |
| SHM2_Seq_int_rev | CCAAATCATCAAAGATGAGGTCG |
| MIS1-1_Seq_ex_fw | GGAGATTTGTTTTCAATGGACC |
| MIS1-1_Seq_ex_rev | GAGATTGGACATGAAAGAAACAGG |
| MIS1-1_Seq_int_fw | CTAGAAATAAAGCAGCTAGCACC |
| MIS1-1_Seq_int_rev | CATTGAGATATCCTGGATGAGTAGG |
| MIS1-2&3_Seq_ex_fw | GTGAGCCTTCAATTACCTCG |
| MIS1-2&3_Seq_ex_rev | GAGATATTGTCAGAATTGTCTTCTGC |
| MIS1-2&3_Seq_int_fw | CGGAACTCCGTTTGTCTATAGCGG |
| MIS1-2&3_Seq_int_rev | CTCCCTTGTCATTGACTTCG |
| BB3rN_Seq_fw | CCACCCCGTAGAAAAGATCAA |
| BB3rN_Seq_rev | CGGCCGTTAAAATACTCA |
| BB3eH_Seq_fw | GAAGCACCGGAAGGAA |
| BB3eH_Seq_rev | ACCTATTCAATGACCAACTCCTGG |
| BB3aK_Seq_fw | GAGGGAGCAGGAGTAGG |
| BB3aK_Seq_rev | AGAAGACCGGTCTTGCTA |

#### 2.3 Sample preparation & GC-TOF-MS analysis of intracellular metabolites

To be able to gather all labelling data of interest by covering a high number of important metabolites, the usage of a diverse set of different GC-MS methods was necessary. Phosphorylated metabolites demand for chemical ionization due to extensive fragmentation in EI, the separation of sugars and their phosphorylated analogous is excellent using ethoximation followed by trimethylsilylation and separation on a 5% phenyl 95% methyl polysiloxane column[4], while specific fragmentation patterns of amino acids can be obtained using electron ionization and preceding tertbutyl-dimethylsilylation. Additionally, the concentration range of intracellular metabolites covers orders of magnitude and split injection as well as splitless injection were necessary for metabolites of high and low concentration, respectively, to avoid to exceed the linear range of the method. Table S5 summarizes, which samples were measured with which GC-MS methods. Table S6 and Table S7 list details for the GC-MS methods and Table S8 lists in detail, which GC-MS method and which data evaluation method was finally used for data evaluation after having applied the selection criteria described in the methods section of the main manuscript.

**Table S5:** Overview of GC-MS methods applied for the analysis of cell extracts of strains grown on differentcarbon sources

| Strain name & carbon source(s) | EtOX/TMS GC-CI-TOFMS, splitless injection | TBDMS GC-EI-TOFMS, splitless injection | TBDMS GC-EI-TOFMS, split1:50 injection |
|---|---|---|---|
| **DasKO MeOH - labelling** | x | x | x |
| **DasKO MeOH+Gly - labelling** | x | x | x |
| **DasKO $CO_2$ - labelling** | x | | x |
| **DasKO FA - labelling** | x | | x |
| **GcvKO MeOH - labelling** | | | x |
| **ShmKO MeOH - labelling** | | | x |
| **Single MisKO MeOH - labelling** | | | x |
| **MisKO MeOH - labelling** | x | | x |
| **MisKO FA - labelling** | x | | x |
| **GcvOE MeOH - labelling** | | | x |
| **Mut- MeOH - labelling** | Zavec et al. [5] | | x |
| **WT FA-labelling** | x | | x |

#### 2.4 GC-TOF-MS data evaluation

**Table S6:** TBDMS GC-EI-TOFMS analytes, retention times, evaluated fragments (fragment structure is explained in more detail by Zamboni et al.[6]) and corresponding isotopologues & m/z ratios

| Metabolite | Name of analyte evaluated * | RT (min) | Ion evaluated ** | Isotopologues | m/z |
|---|---|---|---|---|---|
| Malate | Malate 3TBDMS | 24.37 | $[M\text{-}C_4H_9]^+$ | M+0 | 419.2105 |
| | | | | M+1 | 420.2139 |
| | | | | M+2 | 421.2172 |
| | | | | M+3 | 422.2206 |
| | | | | M+4 | 423.2239 |
| Valine | Valine 2TBDMS | 14.82 | $[M\text{-}C_4H_9]^+$ | M+0 | 288.1815 |
| | | | | M+1 | 289.1849 |
| | | | | M+2 | 290.1882 |
| | | | | M+3 | 291.1916 |
| | | | | M+4 | 292.1949 |
| | | | | M+5 | 293.1983 |
| Glycerate | Glycerate 3TBDMS | 21.61 | $[M\text{-}C_4H_9]^+$ | M+0 | 391.2156 |
| | | | | M+1 | 392.2190 |
| | | | | M+2 | 393.2223 |
| | | | | M+3 | 394.2257 |
| Methionine | Methionine 2TBDMS | 21.65 | $[M\text{-}C_4H_9]^+$ | M+0 | 320.1536 |
| | | | | M+1 | 321.1569 |
| | | | | M+2 | 322.1603 |
| | | | | M+3 | 323.1636 |
| | | | | M+4 | 324.1670 |
| | | | | M+5 | 325.1704 |
| | Methionine DC (decarboxylated***) 2TBDMS | | $[M\text{-}C_5OH_9]^+$ | M+0 | 292.1587 |
| | | | | M+1 | 293.1620 |
| | | | | M+2 | 294.1654 |
| | | | | M+3 | 295.1687 |
| | | | | M+4 | 296.1721 |
| | | | $[M\text{-}C_7O_2SiH_9]^+$ | M+0 | 218.1398 |
| | | | | M+1 | 219.1432 |
| | | | | M+2 | 220.1466 |
| | | | | M+3 | 221.1499 |
| | | | | M+4 | 222.1533 |
| Glutamate | Glutamate 3TBDMS | 26.29 | $[M\text{-}CH_3]^+$ | M+0 | 474.2891 |
| | | | | M+1 | 475.2925 |
| | | | | M+2 | 476.2958 |
| | | | | M+3 | 477.2992 |
| | | | | M+4 | 478.3025 |

| | | | | | |
|---|---|---|---|---|---|
| | | | | M+5 | 479.3059 |
| | | | $[M-C_4H_9]^+$ | M+0 | 432.2422 |
| | | | | M+1 | 433.2455 |
| | | | | M+2 | 434.2489 |
| | | | | M+3 | 435.2522 |
| | | | | M+4 | 436.2556 |
| | | | | M+5 | 437.2589 |
| | Glutamate DC (decarboxylated***) 3TBDMS | | $[M-C_5OH_9]^+$ | M+0 | 404.2472 |
| | | | | M+1 | 405.2506 |
| | | | | M+2 | 406.2540 |
| | | | | M+3 | 407.2573 |
| | | | | M+4 | 408.2607 |
| Alanine | Alanine 2TBDMS | 12.21 | $[M-C_4H_9]^+$ | M+0 | 260.1506 |
| | | | | M+1 | 261.1536 |
| | | | | M+2 | 262.1569 |
| | | | | M+3 | 263.1603 |
| | Alanine DC (decarboxylated***) 2TBDMS | | $[M-C_7O_2SiH_9]^+$ | M+0 | 158.1365 |
| | | | | M+1 | 159.1399 |
| | | | | M+2 | 160.1432 |
| | | | $[M-C_5OH_9]^+$ | M+0 | 232.1553 |
| | | | | M+1 | 233.1586 |
| | | | | M+2 | 234.1620 |
| Glycine | Glycine 2TBDMS | 12.53 | $[M-C_4H_9]^+$ | M+0 | 246.1346 |
| | | | | M+1 | 247.1379 |
| | | | | M+2 | 248.1418 |
| Proline | Proline 2TBDMS | 17.23 | $[M-CH_3]^+$ | M+0 | 328.2128 |
| | | | | M+1 | 329.2162 |
| | | | | M+2 | 330.2195 |
| | | | | M+3 | 331.2229 |
| | | | | M+4 | 332.2262 |
| | | | | M+5 | 333.2296 |
| Serine | Serine 3TBDMS | 22.48 | $[M-C_4H_9]^+$ | M+0 | 390.2316 |
| | | | | M+1 | 391.2350 |
| | | | | M+2 | 392.2383 |
| | | | | M+3 | 393.2417 |
| | | | $[M-CH_3]^+$ | M+0 | 432.2785 |
| | | | | M+1 | 433.2819 |
| | | | | M+2 | 434.2853 |
| | | | | M+3 | 435.2886 |
| | Serine BB (backbone****) | | $[f302]^+$ | M+0 | 302.1976 |
| | | | | M+1 | 303.2005 |

| | | | | | |
|---|---|---|---|---|---|
| | 3TBDMS | | | M+2 | 304.2039 |
| | Serine DC (decarboxylated***) 3TBDMS | | $[M-C_5OH_9]^+$ | M+0 | 362.2367 |
| | | | | M+1 | 363.2400 |
| | | | | M+2 | 364.2434 |
| | | | $[M-C_7O_2SiH_9]^+$ | M+0 | 288.2179 |
| | | | | M+1 | 289.2212 |
| | | | | M+2 | 290.2246 |
| Aspartate | Aspartate 3TBDMS | 24.96 | $[M-C_4H_9]^+$ | M+0 | 418.2265 |
| | | | | M+1 | 419.2299 |
| | | | | M+2 | 420.2332 |
| | | | | M+3 | 421.2366 |
| | | | | M+4 | 422.2399 |
| | | | $[M-CH_3]^+$ | M+0 | 460.2735 |
| | | | | M+1 | 461.2768 |
| | | | | M+2 | 462.2802 |
| | | | | M+3 | 463.2835 |
| | | | | M+4 | 464.2869 |
| | Aspartate BB (backbone****) 3TBDMS | | [f302] $^+$ | M+0 | 302.1976 |
| | | | | M+1 | 303.2005 |
| | | | | M+2 | 304.2039 |
| Lysine | Lysine 3TBDMS | 27.42 | $[M-CH_3]^+$ | M+0 | 473.3415 |
| | | | | M+1 | 474.3448 |
| | | | | M+2 | 475.3482 |
| | | | | M+3 | 476.3515 |
| | | | | M+4 | 477.3549 |
| | | | | M+5 | 478.3583 |
| | | | | M+6 | 479.3616 |
| Fumarate | Fumarate 2TBDMS | 17.40 | $[M-C_4H_9]^+$ | M+0 | 287.1129 |
| | | | | M+1 | 288.1163 |
| | | | | M+2 | 289.1196 |
| | | | | M+3 | 290.1230 |
| | | | | M+4 | 291.1264 |

* the names of the fragments specify the number of tertbutyldimetylsilyl (TBDMS) groups

** the number of the isotopologue denotes the number of $^{13}C$ atoms in the metabolite structure

*** the decarboxylated amino acid corresponds to the derivatized amino acid having lost the C1-carbon atom group[6]

**** the backbone of the amino acids corresponds to the derivatized C1 & C2-carbon atoms only[6], resulting in the fragment with the monoisotopic m/z of 302 ([f302] $^+$)

**Table S7:** EtOx/TMS GC-CI-TOFMS analytes, retention times, evaluated fragments and corresponding isotopologues & m/z ratios

| Metabolite | Name of analyte evaluated * | RT (min) | Ion evaluated** | Isotopologues*** | m/z |
|---|---|---|---|---|---|
| Aspartate | Aspartate 2TMS | 12.43 | $[M+H]^+$ | M+0 | 278.1238 |
| | | | | M+1 | 279.1272 |
| | | | | M+2 | 280.1305 |
| | | | | M+3 | 281.1338 |
| | | | | M+4 | 282.1373 |
| R5P | Ribose 5-phosphate 5TMS 1EtOx | 24.29 | $[M-CH_3]^+$ | M+0 | 618.2350 |
| | | | | M+1 | 619.2383 |
| | | | | M+2 | 620.2417 |
| | | | | M+3 | 621.2450 |
| | | | | M+4 | 622.2484 |
| | | | | M+5 | 623.2517 |
| S7P | Sedoheptulose 7-phosphate 7TMS 1EtOx | 32.26 | $[M-CH_3]^+$ | M+0 | 822.3351 |
| | | | | M+1 | 823.3385 |
| | | | | M+2 | 824.3419 |
| | | | | M+3 | 825.3452 |
| | | | | M+4 | 826.3486 |
| | | | | M+5 | 827.3519 |
| | | | | M+6 | 828.3553 |
| | | | | M+7 | 829.3586 |
| PEP | Phosphoenol-pyruvate 3TMS | 14.49 | $[M+H]^+$ | M+0 | 385.1088 |
| | | | | M+1 | 386.1121 |
| | | | | M+2 | 387.1155 |
| | | | | M+3 | 388.1188 |
| | | | $[M-CH_3]^+$ | M+0 | 369.0775 |
| | | | | M+1 | 370.0808 |
| | | | | M+2 | 371.0842 |
| | | | | M+3 | 372.0875 |
| 2-PG | 2-Phospho-glycerate 4TMS | 17.10 | $[M+H]^+$ | M+0 | 475.1583 |
| | | | | M+1 | 476.1617 |
| | | | | M+2 | 477.1650 |
| | | | | M+3 | 478.1684 |

| | | | | | |
|---|---|---|---|---|---|
| 3-PG | 3-Phospho-glycerate 4TMS | 17.51 | $[M+H]^+$ | M+0 | 475.1583 |
| | | | | M+1 | 476.1617 |
| | | | | M+2 | 477.1650 |
| | | | | M+3 | 478.1684 |
| AKG | α-Ketoglutarate 2TMS 1EtOx | 14.61 | $[M+H]^+$ | M+0 | 334.1506 |
| | | | | M+1 | 335.154 |
| | | | | M+2 | 336.1573 |
| | | | | M+3 | 337.1607 |
| | | | | M+4 | 338.164 |
| | | | | M+5 | 339.1674 |
| | | | $[M-CH_3]^+$ | M+0 | 318.1193 |
| | | | | M+1 | 319.1210 |
| | | | | M+2 | 320.1186 |
| | | | | M+3 | 321.1200 |
| | | | | M+4 | 322.1181 |
| | | | | M+5 | 323.1163 |
| I-Cit | Iso-citrate 4TMS | 17.76 | $[M+H]^+$ | M+0 | 481.1924 |
| | | | | M+1 | 482.1957 |
| | | | | M+2 | 483.1991 |
| | | | | M+3 | 484.2025 |
| | | | | M+4 | 485.2058 |
| | | | | M+5 | 486.2092 |
| | | | | M+6 | 487.2125 |
| | | | $[M-CH_3]^+$ | M+0 | 465.1616 |
| | | | | M+1 | 466.165 |
| | | | | M+2 | 467.1683 |
| | | | | M+3 | 468.1717 |
| | | | | M+4 | 469.1751 |
| | | | | M+5 | 470.1784 |
| | | | | M+6 | 471.1818 |
| G6P | Glucose 6-phosphate 6TMS 1EtOx | 28.85 | $[M-CH_3]^+$ | M+0 | 720.2851 |
| | | | | M+1 | 721.2884 |
| | | | | M+2 | 722.2918 |
| | | | | M+3 | 723.2951 |
| | | | | M+4 | 724.2985 |

| | | | | | |
|---|---|---|---|---|---|
| | | | | M+5 | 725.3018 |
| | | | | M+6 | 726.3052 |
| F6P | Fructose 6-phosphate 6TMS 1EtOx | 28.53 | $[M-CH_3]^+$ | M+0 | 720.2851 |
| | | | | M+1 | 721.2884 |
| | | | | M+2 | 722.2918 |
| | | | | M+3 | 723.2951 |
| | | | | M+4 | 724.2985 |
| | | | | M+5 | 725.3018 |
| | | | | M+6 | 726.3052 |

* the names of the fragments specify the number of trimethylsilyl (TMS) and ethoxyamino (EtOx) groups

** the number of the isotopologue denotes the number of $^{13}C$ atoms in the metabolite structure

**Table S8:** List of GC-MS & data evaluation methods and samples they were applied to

| Metabolite / fragment name | GC-MS method | | Data evaluation method | | | Applied for the following samples* |
|---|---|---|---|---|---|---|
| | Derivatization / Ioniziation | injection | Fragment/ Adduct | MS data type | Mass ex-traction window [±ppm] | |
| Malate | TBDMS GC-EI-TOFMS | split 1:50 | $[M-C_4H_9]^+$ | profile | 50 | DasKO MeOH, DasKO MeOH+Gly, DasKO reverse labelling |
| | TBDMS GC-EI-TOFMS | split 1:50 | $[M-C_4H_9]^+$ | centroid | 50 | GcvKO MeOH, ShmKO MeOH, Single MisKO MeOH, MisKO MeOH, GcvOE MeOH, DasKO FA, WT FA, MisKO FA |
| | EtOx/TMS GC-CI-TOFMS | splitless | $[M+H]^+$ | profile | 50 | Mut(-) MeOH** |
| Valine | TBDMS GC-EI-TOFMS | splitless | $[M-C_4H_9]^+$ | profile | 50 | DasKO MeOH, DasKO MeOH+Gly, |
| | TBDMS GC-EI-TOFMS | split 1:50 | $[M-C_4H_9]^+$ | profile | 50 | DasKO reverse labelling, GcvKO MeOH, GcvOE MeOH |
| | EtOx/TMS GC-CI-TOFMS | splitless | $[M-CH_3]^+$ | profile | 15 | Mut(-) MeOH** |
| Glycerate | TBDMS GC-EI-TOFMS | splitless | $[M-C_4H_9]^+$ | profile | 50 | DasKO MeOH, DasKO MeOH+Gly |
| Methionine | TBDMS GC-EI-TOFMS | splitless | $[M-C_4H_9]^+$ | profile | 50 | DasKO MeOH 72h, DasKO MeOH+Gly 72h |
| | TBDMS GC-EI-TOFMS | splitless | $[M-C_4H_9]^+$ | centroid | 50 | DasKO MeOH 2h 24h, DasKO MeOH+Gly 2h 24h |
| | TBDMS GC-EI-TOFMS | split 1:50 | $[M-C_4H_9]^+$ | profile | 50 | DasKO reverse labelling, GcvKO MeOH, GcvOE OE MeOH |
| | TBDMS GC-EI-TOFMS | split 1:50 | $[M-C_4H_9]^+$ | centroid | 50 | Mut(-) MeOH, ShmKO MeOH, Single MisKO MeOH, MisKO MeOH, WT FA, DasKO FA, MisKO FA |
| Methionine DC | TBDMS GC-EI-TOFMS | splitless | $[M-C_5OH_9]^+$ | profile | 50 | DasKO MeOH 2h 72h, DasKO MeOH+Gly 2h 72h |
| | TBDMS GC-EI-TOFMS | splitless | $[M-C_7O_2SiH_9]^+$ | profile | 50 | DasKO MeOH 24h, DasKO MeOH+Gly 24h |

| | | | | | | |
|---|---|---|---|---|---|---|
| | TBDMS GC-EI-TOFMS | split 1:50 | $[M-C_5OH_9]^+$ | profile | 50 | DasKO reverse labelling 0h 72h, ShmKO MeOH, Mut(-) MeOH |
| | TBDMS GC-EI-TOFMS | split 1:50 | $[M-C_7O_2SiH_9]^+$ | profile | 50 | DasKO reverse labelling 24h, GcvKO MeOH, GcvOE MeOH |
| | TBDMS GC-EI-TOFMS | split 1:50 | $[M-C_5OH_9]^+$ | centroid | 50 | MisKO MeOH, WT FA, DasKO FA, MisKO FA |
| Glutamate | TBDMS GC-EI-TOFMS | splitless | $[M-CH_3]^+$ | profile | 50 | DasKO MeOH 2h 72h, DasKO MeOH+Gly 2h 72h |
| | TBDMS GC-EI-TOFMS | split 1:50 | $[M-CH_3]^+$ | profile | 50 | DasKO MeOH 24h, DasKO reverse labelling |
| | TBDMS GC-EI-TOFMS | split 1:50 | $[M-C_4H_9]^+$ | centroid | 50 | MisKO MeOH, DasKO FA |
| | TBDMS GC-EI-TOFMS | split 1:50 | $[M-CH_3]^+$ | centroid | 50 | MisKO FA, WT FA |
| Glutamate DC | TBDMS GC-EI-TOFMS | split 1:50 | $[M-C_5OH_9]^+$ | centroid | 50 | MisKO MeOH, MisKO FA, WT FA, DasKO FA |
| Alanine | TBDMS GC-EI-TOFMS | splitless | $[M-C_4H_9]^+$ | profile | 50 | DasKO MeOH, DasKO MeOH+Gly |
| | TBDMS GC-EI-TOFMS | split 1:50 | $[M-C_4H_9]^+$ | profile | 50 | DasKO reverse labelling |
| | TBDMS GC-EI-TOFMS | split 1:50 | $[M-C_4H_9]^+$ | centroid | 50 | MisKO MeOH, WT FA, DasKO FA |
| Alanine DC | TBDMS GC-EI-TOFMS | split 1:50 | $[M-C_7O_2SiH_9]^+$ | profile | 50 | DasKO MeOH, DasKO MeOH+Gly |
| | TBDMS GC-EI-TOFMS | split 1:50 | $[M-C_5OH_9]^+$ | centroid | 50 | MisKO MeOH, DasKO FA, WT FA, MisKO FA |
| Glycine | TBDMS GC-EI-TOFMS | split 1:50 | $[M-C_4H_9]^+$ | profile | 50 | DasKO MeOH, DasKO MeOH+Gly, DasKO reverse labelling, GcvKO MeOH, ShmKO MeOH, single MisKO MeOH, GcvOE MeOH, DasKO FA 24h |
| | TBDMS GC-EI-TOFMS | split 1:50 | $[M-C_4H_9]^+$ | centroid | 50 | MisKO MeOH, WT FA, MisKO FA, DasKO FA 2h 72h |
| | EtOx/TMS GC-CI-TOFMS | splitless | $[M-CH_3]^+$ | profile | 50 | Mut(-) MeOH** |
| Proline | TBDMS GC-EI-TOFMS | split 1:50 | $[M-CH_3]^+$ | profile | 50 | DasKO MeOH 24h 72h, DasKO MeOH+Gly |

| | | | | | | |
|---|---|---|---|---|---|---|
| Serine | TBDMS GC-EI-TOFMS | split 1:50 | $[M-C_4H_9]^+$ | profile | 50 | DasKO MeOH, DasKO MeOH+Gly, DasKO reverse labelling, GcvKO MeOH, ShmKO MeOH, single MisKO MeOH, GcvOE MeOH, Mut(-) MeOH |
| | TBDMS GC-EI-TOFMS | split 1:50 | $[M-C_4H_9]^+$ | centroid | 50 | MisKO MeOH, WT FA, DasKO FA |
| | TBDMS GC-EI-TOFMS | split 1:50 | $[M-CH_3]^+$ | centroid | 50 | MisKO FA |
| Serine BB | TBDMS GC-EI-TOFMS | split 1:50 | $[f302]^+$ | profile | 50 | DasKO MeOH, DasKO MeOH+Gly, DasKO reverse labelling, GcvKO MeOH, ShmKO MeOH, single MisKO MeOH, GcvOE MeOH, Mut(-) MeOH, DasKO FA |
| | TBDMS GC-EI-TOFMS | split 1:50 | $[f302]^+$ | centroid | 50 | MisKO MeOH, MisKO FA, WT FA |
| Serine DC | TBDMS GC-EI-TOFMS | split 1:50 | $[M-C_7O_2SiH_9]^+$ | profile | 50 | DasKO MeOH 2h 24h, DasKO MeOH+Gly 2h 24h, DasKO reverse labelling 2h 24h, ShmKO MeOH, GcvKO MeOH, GcvOE MeOH, Mut(-) MeOH |
| | TBDMS GC-EI-TOFMS | split 1:50 | $[M-C_5OH_9]^+$ | profile | 50 | DasKO MeOH 72h, DasKO MeOH+Gly 72h, DasKO reverse labelling 72h |
| | TBDMS GC-EI-TOFMS | split 1:50 | $[M-C_7O_2SiH_9]^+$ | centroid | 50 | MisKO MeOH, MisKO FA, WT FA, DasKO FA |
| Aspartate | EtOx/TMS GC-CI-TOFMS | splitless | $[M+H]^+$ | profile | 50 | DasKO MeOH 72h, DasKO MeOH+Gly 72h |
| | TBDMS GC-EI-TOFMS | split 1:50 | $[M-C_4H_9]^+$ | profile | 50 | DasKO MeOH 2h 24h, DasKO MeOH+Gly 2h 24h |
| | TBDMS GC-EI-TOFMS | split 1:50 | $[M-CH_3]^+$ | profile | 50 | DasKO reverse labelling, GcvKO MeOH, ShmKO, single MisKO MeOH, GcvOE MeOH, Mut(-) MeOH, WT FA 2h |
| | TBDMS GC-EI-TOFMS | split 1:50 | $[M-CH_3]^+$ | centroid | 50 | MisKO MeOH, MisKO FA, DasKO FA, WT FA 24h 72h |

| | | | | | | |
|---|---|---|---|---|---|---|
| Aspartate BB | TBDMS GC-EI-TOFMS | split 1:50 | $[f302]^+$ | profile | 50 | DasKO MeOH, DasKO MeOH+Gly |
| | TBDMS GC-EI-TOFMS | split 1:50 | $[f302]^+$ | centroid | 50 | DasKO FA 2h 24h, WT FA 2h 24h |
| Lysine | TBDMS GC-EI-TOFMS | split 1:50 | $[M-CH_3]^+$ | profile | 50 | DasKO MeOH 24h 72h, DasKO MeOH+Gly 24h 72h, DasKO reverse labelling 24h 72h |
| | EtOx/TMS GC-CI-TOFMS | splitless | $[M-CH_3]^+$ | profile | 50 | Mut(-) MeOH** |
| R5P | EtOx/TMS GC-CI-TOFMS | splitless | $[M-CH_3]^+$ | profile | 50 | DasKO MeOH, DasKO MeOH+Gly, WT FA 2h, Mut(-) MeOH ** |
| | EtOx/TMS GC-CI-TOFMS | splitless | $[M-CH_3]^+$ | centroid | 50 | DasKO FA, WT FA 24h 72h |
| S7P | EtOx/TMS GC-CI-TOFMS | splitless | $[M-CH_3]^+$ | profile | 50 | DasKO MeOH, DasKO MeOH+Gly, DasKO FA 2h 24h, Mut(-) MeOH ** |
| | EtOx/TMS GC-CI-TOFMS | splitless | $[M-CH_3]^+$ | centroid | 50 | WT FA, DasKO FA 72h |
| PEP | EtOx/TMS GC-CI-TOFMS | splitless | $[M+H]^+$ | profile | 15 | DasKO MeOH 24h 72h, DasKO MeOH+Gly 24h 72h, MisKO MeOH |
| | EtOx/TMS GC-CI-TOFMS | splitless | $[M-CH_3]^+$ | profile | 15 | DasKO MeOH 2h, DasKO MeOH+Gly 2h, DasKO FA 72h |
| | EtOx/TMS GC-CI-TOFMS | splitless | $[M+H]^+$ | centroid | 15 | MisKO MeOH, MisKO FA, WT FA 24h |
| | EtOx/TMS GC-CI-TOFMS | splitless | $[M-CH_3]^+$ | centroid | 15 | WT FA 72h |
| 2-PG | EtOx/TMS GC-CI-TOFMS | splitless | $[M+H]^+$ | profile | 50 | DasKO MeOH, DasKO MeOH+Gly, DasKO FA 24h, WT FA 24h, Mut(-) MeOH ** |
| | EtOx/TMS GC-CI-TOFMS | splitless | $[M+H]^+$ | centroid | 50 | WT FA 72h |
| | EtOx/TMS GC-CI-TOFMS | splitless | $[M+H]^+$ | profile | 15 | DasKO FA 72h |
| 3-PG | EtOx/TMS GC-CI-TOFMS | splitless | $[M+H]^+$ | profile | 50 | DasKO MeOH, DasKO MeOH+Gly, Mut(-) MeOH **, WT FA 2h |
| | EtOx/TMS GC-CI-TOFMS | splitless | $[M+H]^+$ | centroid | 50 | MisKO MeOH, DasKO FA, WT FA 24h 72h |
| AKG | EtOx/TMS GC-CI-TOFMS | splitless | $[M+H]^+$ | centroid | 50 | DasKO MeOH 24h |

| | | | | | | |
|---|---|---|---|---|---|---|
| | EtOx/TMS GC-CI-TOFMS | splitless | $[M\text{-}CH_3]^+$ | centroid | 50 | DasKO MeOH 72h, WT FA |
| | EtOx/TMS GC-CI-TOFMS | splitless | $[M+H]^+$ | profile | 50 | DasKO MeOH 2h |
| | EtOx/TMS GC-CI-TOFMS | splitless | $[M\text{-}CH_3]^+$ | profile | 50 | DasKO FA |
| I-Cit | EtOx/TMS GC-CI-TOFMS | splitless | $[M\text{-}CH_3]^+$ | centroid | 50 | DasKO MeOH 72h, WT FA 2h 24h |
| | EtOx/TMS GC-CI-TOFMS | splitless | $[M+H]^+$ | centroid | 15 | DasKO MeOH 2h 24h, Mut(-) MeOH **, DasKO reverse labelling |
| | EtOx/TMS GC-CI-TOFMS | splitless | $[M\text{-}CH_3]^+$ | centroid | 50 | MisKO FA |
| Fumarate | TBDMS GC-EI-TOFMS | s split 1:50 | $[M\text{-}C_4H_9]^+$ | centroid | 50 | DasKO MeOH, DaSKO MeOH+Gly, DasKO reverse labelling, GcvKO MeOH, GcvOE MeOH, MisKO MeOH, MisKO FA, DasKO FA, WT FA |
| G6P | EtOx/TMS GC-CI-TOFMS | splitless | $[M\text{-}CH_3]^+$ | profile | 50 | DasKO MeOH 2h 24h, DasKO MeOH+Gly 2h 24h |
| | EtOx/TMS GC-CI-TOFMS | splitless | $[M\text{-}CH_3]^+$ | centroid | 50 | DasKO FA 2h 24h, |
| F6P | EtOx/TMS GC-CI-TOFMS | splitless | $[M\text{-}CH_3]^+$ | centroid | 50 | WT FA |

*if no time point is specified, all time points of the strain and carbon source were evaluated by the respective method

** GC-MS and data evaluation method as described in Zavec et al. [5]
